# Integrated Genome and Transcriptome Analyses Reveal the Mechanism of Genome Instability in Ataxia with Oculomotor Apraxia 2

**DOI:** 10.1101/2021.05.07.443085

**Authors:** Radhakrishnan Kanagaraj, Richard Mitter, Theodoros Kantidakis, Matthew M. Edwards, Anaid Benitez, Probir Chakravarty, Beiyuan Fu, Olivier Becherel, Fengtang Yang, Martin F. Lavin, Amnon Koren, Aengus Stewart, Stephen C. West

## Abstract

Mutations in the *SETX* gene, which encodes Senataxin, are associated with the progressive neurodegenerative diseases Ataxia with Oculomotor Apraxia 2 (AOA2) and Amyotrophic Lateral Sclerosis 4 (ALS4). To identify the causal defect in AOA2, patient-derived cells and *SETX* knockouts (human and mouse) were analyzed using integrated genomic and transcriptomic approaches. We observed a genome-wide increase in chromosome instability (gains and losses) within genes and at chromosome fragile sites, resulting in changes to gene expression profiles. Senataxin loss caused increased transcription stress near promoters that correlated with high GCskew and R-loop accumulation at promoter-proximal regions. Notably, the chromosomal regions with gains and losses overlapped with regions of elevated transcription stress. In the absence of Senataxin, we found that Cockayne Syndrome protein CSB was required for the recruitment of the transcription-coupled repair endonucleases (XPG and XPF) and recombination protein RAD52 to target and resolve transcription bubbles containing R-loops, leading to error prone repair and genomic instability. These results show that transcription stress is an important contributor to *SETX* mutation-associated chromosome fragility and AOA2.

## INTRODUCTION

Transcription has been linked to mutagenesis, DNA breakage and genomic instability (1). Recent studies have highlighted the consequences of transcription-replication conflicts and the formation of transcription-linked R-loops as sources of genomic instability in both prokaryotes and eukaryotes (2–5). R-loops are three-stranded nucleic acid structures containing an RNA/DNA hybrid and an unpaired single-strand of DNA. They are found near gene promoters and terminators (6–9), rDNA repeats (10), tRNA genes (11), DNA double-strand breaks (DSBs) (12), replication origins (13, 14) and immunoglobulin class-switch regions (15).

R-loops are thought to have physiological functions, which include regulating gene expression, facilitating transcription termination and promoting class switch recombination (4,8,15,16). However, aberrant R-loop formation and improper processing of these structures contributes to hypermutation, DSB formation, and chromosome rearrangements, which are all sources of genomic instability and human disease (4,5,17). The proper regulation of R-loop homeostasis is therefore vital for the maintenance of genome integrity.

Eukaryotic cells have evolved multiple mechanisms to control R-loop formation. Unscheduled or unwanted R-loops are either degraded by the ribonucleases RNaseH1 and RNaseH2, or removed by RNA/DNA helicases such as Senataxin (Sen1 in yeast), Aquarius or UAP56 (9,18–22). Senataxin (SETX) was first identified due to its association with an inherited autosomal recessive adolescent onset disorder known as Ataxia with Oculomotor Apraxia 2 (AOA2) (23). Mutations in the *SETX* gene are also linked to a rare, dominantly inherited, form of motor neuron disease Amyotrophic Lateral Sclerosis 4 (ALS4) (24). *SETX* mutations associated with AOA2 and ALS4 are generally considered to be loss-of-function and gain-of-function, respectively (25). AOA2 is characterized by cerebellar atrophy, early loss of reflexes, late peripheral neuropathy, oculomotor apraxia and impaired motor functions (26). Patient-derived AOA2 cells are sensitive to DNA damaging agents including H_2_O_2_ (27–29). AOA2 cells exhibit altered gene expression (including neuronal genes) and increased R-loop levels (30). Although a *Setx* knockout mouse has been generated, it fails to exhibit the neurodegenerative features typical of afflicted individuals (31). However, the male mice were infertile and SETX was shown to be essential for the removal of R-loops during meiotic recombination in spermatocytes.

Senataxin has been implicated in the resolution of R-loops that form during transcription regulation (32), transcription termination (9,33–35), replication-transcription collisions (36, 37), DNA damage (38–40), meiotic gene silencing (41) and the antiviral transcriptional response (42). However, the precise molecular functions of *SETX,* and how mutations in this gene lead to AOA2 neuropathy, remain largely unknown.

In this study, we provide the first detailed genome-wide analysis of cells derived from AOA2 patients and *SETX* knockouts (human and mouse). Using a variety of genomic and transcriptomic methods, we show that loss of SETX leads to a genome-wide increase in RNAPII levels (transcription stress) and chromosome instability across genes and at fragile sites. Importantly, transcription stress near promoters correlated with high GCskew and R-loop accumulation at promoter-proximal regions. In the absence of SETX, R-loops are targeted and repaired by transcription-coupled repair (TCR) and recombination proteins. The reactions are error prone and lead to genomic instability that are the likely cause of neurodegeneration.

## RESULTS

### AOA2 cells exhibit transcription-dependent genome instability

To investigate the genome-wide chromosome instability/fragility phenotypes associated with SETX-deficiency, we analysed an AOA2 fibroblast cell line (designated AOA2-P1) that has a large deletion (exons 16-23) in the helicase domain of SETX (Figure 1A) (29). Immunostaining for the DNA damage response protein 53BP1 revealed a four-fold increase in the number of 53BP1 nuclear bodies (NBs) in Cyclin A-negative G1 cells compared to control (CTRL-C1) fibroblasts, which was suppressed by treatment with the transcription inhibitor cordycepin (Figure 1B and C). The AOA2-P1 cells also showed a ∼5-fold increase in the formation of micronuclei compared with control cells, which was again suppressed by cordycepin treatment (Figure 1D and E). 53BP1 NB formation and micronuclei were also visualised by live cell imaging of U2OS cells that stably expressed GFP-tagged 53BP1, following treatment with control (Supplementary Movie S1) or *SETX* siRNA (Supplementary Movie S2).

**Figure 1.**
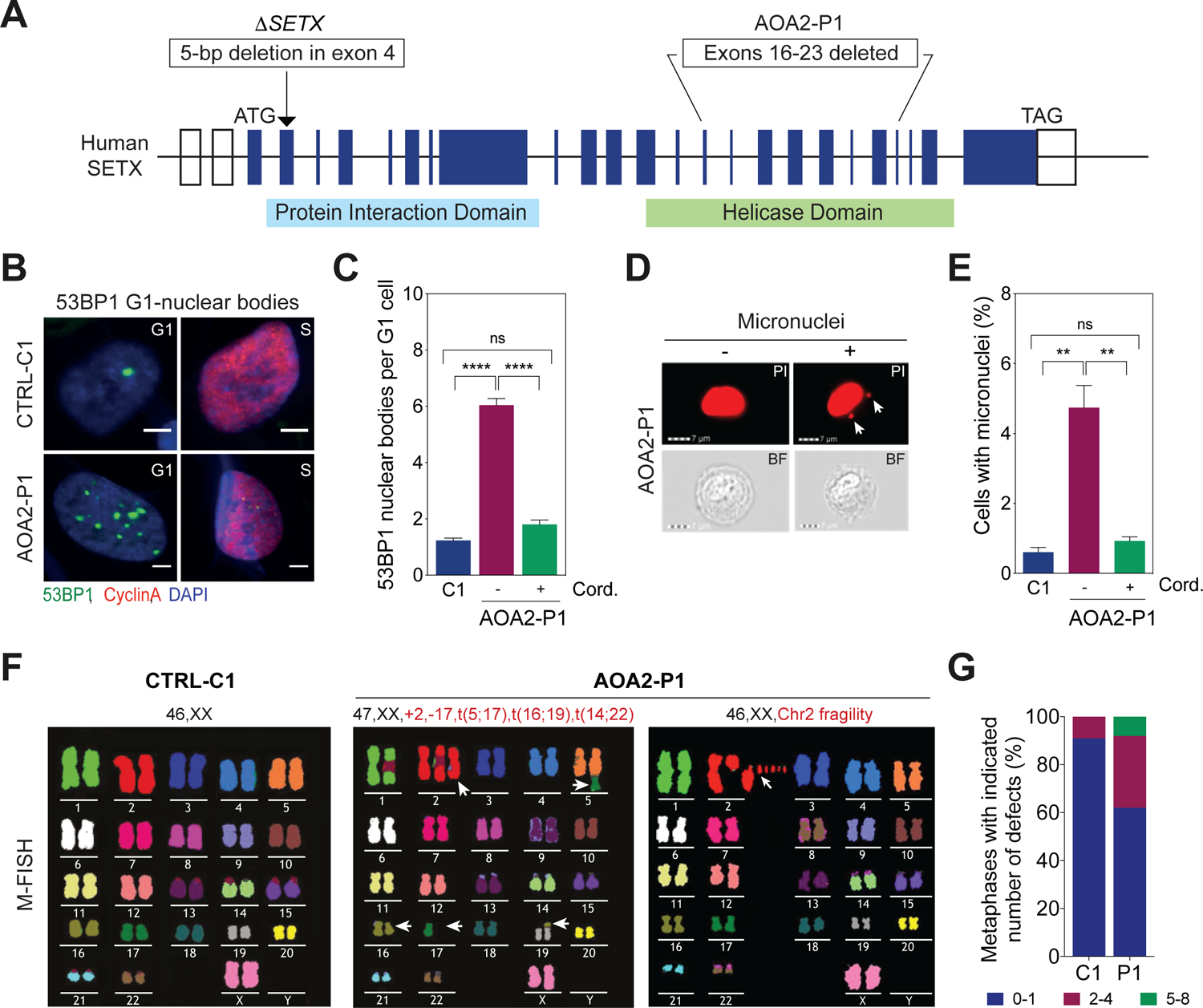
SETX deficiency promotes chromosome fragility. (A) Diagram of human *SETX* showing Δ*SETX* and the AOA2-P1 mutant fibroblasts. Boxes representing exons and the positions of start (ATG) and stop (TAG) codons are shown. Protein interaction and helicase domains are indicated. (B) Immunostaining with 53BP1 (green) and Cyclin A (red) antibodies in control (CTRL-C1) and AOA2-P1 fibroblasts. DNA was stained with DAPI (blue). Representative images of G1- and S-phase cells are shown. Scale bar, 5μm. (C) Quantification of 53BP1 nuclear bodies in G1 cells, as in (B), treated with or without cordycepin (Cord.). Data represent the mean ± s.e.m of three independent experiments with >300 cells per condition. (D) Imagestream imaging flow cytometry of AOA2-P1 fibroblasts. Nuclei were stained with propidium iodide (PI). Representative PI and bright field (BF) images of mononucleated cells with and without micronuclei are shown. White arrows denote micronuclei. Scale bar, 7 μm. (E) Quantification of cells with micronuclei, as in (D). Cells were treated with or without cordycepin. The data represent the mean ± s.e.m of three independent experiments with >4000 cells per condition. (F) Multicolour fluorescent *in situ* hybridization (M-FISH) analyses of metaphase spreads from CTRL-C1 and two AOA2-P1 fibroblasts showing deletions/amplifications and chromosome fragility. Representative karyotypes are shown. White arrows indicate chromosome aberrations. (G) Quantification of metaphases with indicated number of aberrations, as in (F). 30 metaphases were analysed per condition. **P<0.01 and ****P<0.0001 by Mann-Whitney test. P≥0.05 is considered not significant (ns).

Analysis of AOA2-P1 fibroblasts by multiplex fluorescence *in situ* hybridization (M-FISH) and DAPI-banding revealed extensive chromosomal rearrangements (deletions, amplifications and translocations) and numerical aberrations (Figure 1F and Supplementary Figure S1A). We found that 38% of the AOA2-P1 cells exhibited two or more aberrations compared to 9% in control cells (Figure 1G). Similar results were observed with human HAP1-*SETX* knockout cells (Δ*SETX*; Figure S1B) made by CRISPR/Cas9-mediated gene targeting (Figure S1C-G). These results show that loss or inactivation of SETX leads to transcription-dependent genomic instability.

### SETX deficiency induces genome-wide copy number changes

Copy number changes (CNCs), such as submicroscopic deletions (losses) and amplifications (gains), can be caused by transcription-dependent genomic instability, as identified using array comparative genomic hybridization (aCGH) (43, 44). To determine whether SETX-deficient cells exhibit CNCs, genomic DNAs extracted from AOA2-P1 and CTRL-C1 cells were compared by aCGH using whole-genome arrays containing 400K unique oligonucleotide sequences. We observed 57 and 48 CNCs in two independent experiments (Figure 2A and Supplementary Figure S2A), and of these 47 CNCs (23 gains and 24 losses) were common to both experiments (Figure 2B). Remarkably, 63% of the CNCs located to regions containing known genes, indicating their association with transcription (Supplementary Table S1A and B). Permutation-based overlap comparisons of CNCs and previously identified fragile sites revealed that the gains, but not losses, were enriched for early replicating fragile sites (ERFS) (45), which colocalize with highly expressed gene clusters (Figure 2C and Supplementary Figure S2B). We also found that gain regions were significantly enriched for genomic locations of double-strand breaks (DSBs) induced by neocarzinostatin (NCS-Breakome), but not those induced by aphidicolin (APH-Breakome) (46) (Figure 2D, 2E and Supplemenary Figure S2B). Notably, 15 gains and 11 losses overlapped with common/rare fragile sites (C/RFS) (47), and many of the genes in these regions mapped to common fragile sites (CFSs) (48) such as *CDH13*, *PARK2* and *WWOX* (Supplementary Table S1). We were, however, unable to confirm the statistical significance of this overlap with permutation-based tests, likely because CFSs and C/RFSs cover large portions of the genome (Supplementary Figure S2C-E).

**Figure 2.**
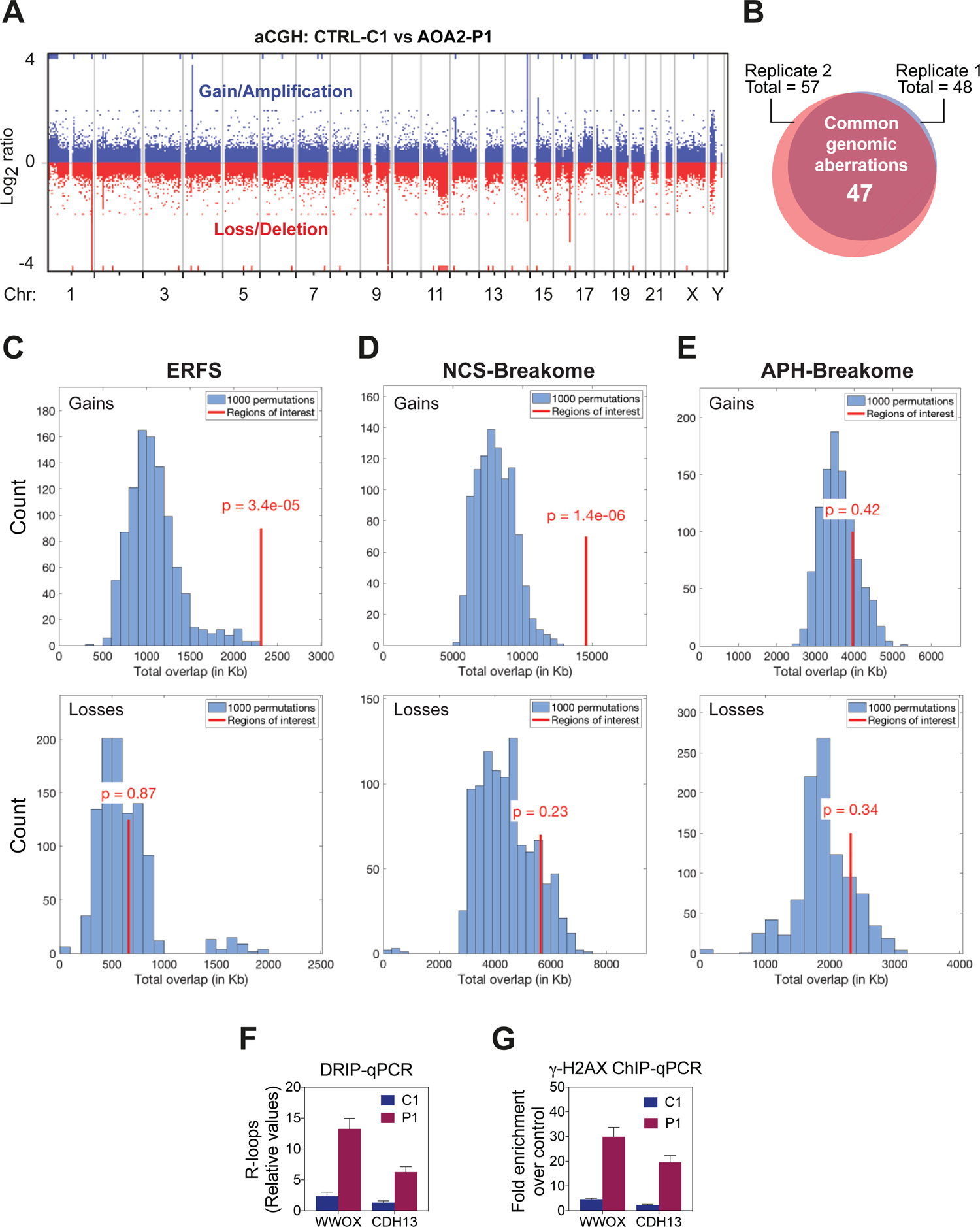
Genome-wide chromosome instability in AOA2 fibroblasts. (A) Array comparative genomic hybridization (aCGH) analyses were carried out with genomic DNA from CTRL-C1 and AOA2-P1 fibroblasts. Representative whole genome plot of aCGH profile is shown. Blue and red lines indicate chromosome regions with gains/amplifications and losses/deletions, respectively. (B) Venn diagram showing the overlap of chromosome aberrations detected in two independent aCGH experiments (replicates 1 and 2), as in (A). (C) Upper panel: Histogram showing overlaps between 1000 permuted AOA2-P1 fibroblast gain regions with early replicating fragile sites (ERFS). Lower panel: Histogram showing overlaps between 1000 permuted AOA2-P1 fibroblast loss regions with the ERFSs. Red line indicates the degree of overlap (in kb). P values for the overlap compared to permutations are shown, with P<0.05 considered a significant enrichment/depletion. (D and E) As (C) showing the overlaps between AOA2-P1 fibroblast gain and loss regions with neocarzinostatin-breakome and aphidicolin-breakome sensitive regions. (F) DRIP-qPCR analyses, using the R-loop specific S9.6 monoclonal antibody, were carried out with genomic DNA from CTRL-C1 and AOA2-P1 fibroblasts. R-loop prone regions at *WWOX* and *CDH13* genes were analysed. Relative values of R-loops immunoprecipitated in each region, normalized to input values and to the signal at the SNRPN negative control region are shown. Data represent the mean ± s.e.m of three independent experiments. (G) ChIP-qPCR analyses at *WWOX* and *CDH13* genes using a γ-H2AX antibody was carried out with cross-linked chromatin from CTRL-C1 and AOA2-P1 fibroblasts. Fold enrichment was calculated as a ratio of γ-H2AX antibody signal versus control IgG. Data represent the mean ± s.e.m of three independent experiments.

CFSs represent unusually long genes that are difficult to transcribe and/or replicate, where collisions between the replisome and transcription apparatus lead to R-loop formation and DNA breakage. We therefore used DNA-RNA immunoprecipitation (DRIP) to investigate the accumulation of R-loops at two CFS loci (*WWOX* and *CDH13*) in genomic DNA from the AOA2-P1 and CTRL-C1 fibroblasts. Quantitative real-time PCR (qPCR) analyses revealed increased R-loop formation at both loci in AOA2-P1 compared to the CTRL-C1 (Figure 2F). Moreover, these CFS loci accumulated significant levels of DNA damage, as measured by γ-H2AX chromatin immunoprecipitation (ChIP) (Figure 2G).

Τo confirm that loss of SETX induces genome-wide CNCs, aCGH was used to compare genomic DNA from five different patient-derived AOA2 lymphoblastoid cell lines (LCLs; P1 to P4 and P1.1; Figure 3A) with their respective controls (C1 to C4 and C1.1). A significant number of CNCs (369 gains and losses) were identified (Figure 3B, Supplementary Figure S3A, and Supplementary Table S2A-J), and mapped mainly to gene-rich regions (81%; Figure 3B, 3D and 3E). Overlap analyses revealed that both gain and loss regions were significantly enriched for GC-rich and transcriptionally-active ERFS (P=8 × 10^−06^ and P=1.9 × 10^−02^, respectively, by permutation test), and only loss regions were significantly enriched for NCS-breakome sensitive regions (NSR) of the genome (P=4.3 × 10^−03^ by permutation test) (Figure 3B). Overlap tests revealed no significant enrichment of the APH-breakome sensitive regions (ASR). Consistent with our findings in AOA2-P1 fibroblasts, we identified 78 gains and 81 losses that overlapped with the C/RFSs, and 34 gains and 37 losses that overlapped with CFSs.

**Figure 3.**
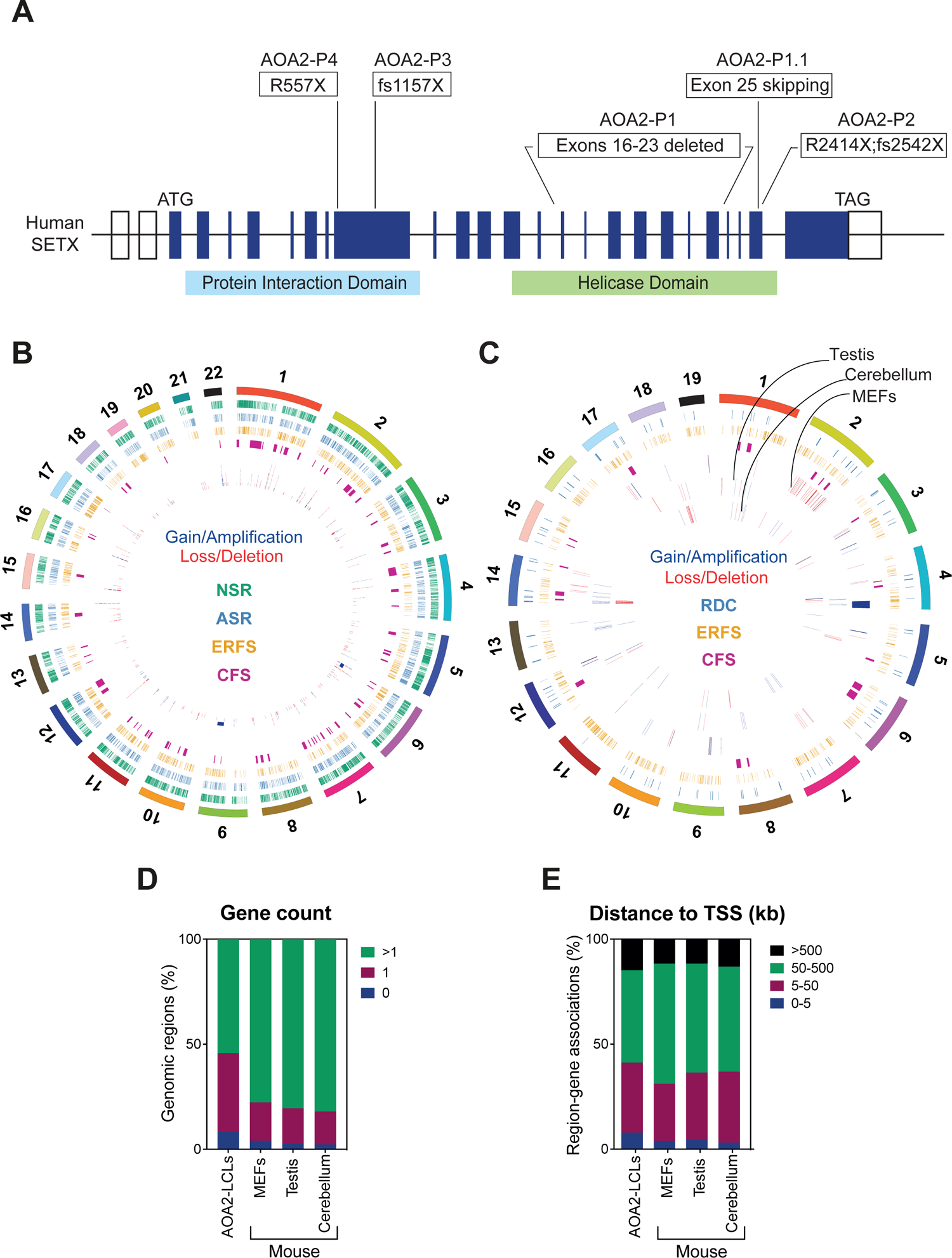
Hotspots of genomic instability in SETX-deficient human and mouse cells. (A) Diagram of human *SETX*, as in Figure 1A, showing the mutations in the AOA2 LCLs. (B) Circos plot depicting the identified CNCs in AOA2 LCLs. Gains/amplifications and losses/deletions are marked in blue and red, respectively. Chromosome locations of known fragile sites are marked in magenta (CFS), orange (ERFS), cyan (ASR) and green (NSR). The outer ring shows all chromosomes. CFS, common fragile site; ERFS, early replicating fragile site; ASR, aphidicolin-breakome sensitive region; NSR, neocarzinostatin-breakome sensitive region. (C) As (B) except that the Circos plot depicts CNCs identified in MEFs, testis and cerebellum of *Setx*^-/-^ mice. RDC, recurrent DNA break cluster. (D) Genomic Regions Enrichment of Annotations Tool (GREAT) analyses were carried out with aCGH datasets from AOA2-LCLs, mouse cells and tissues (MEFs, testis and cerebellum). Plot depicts the percentage of genomic regions associated with the indicated gene count. (E) Plot depicting the distance of genomic regions to their nearest transcription start sites (TSSs), identified in AOA2 and mouse aCGH experiments. Distances are indicated in kilobases (kb).

To confirm and extend these results, aCGH experiments were performed using genomic DNA isolated from MEFs, testis and cerebellar vermis isolated from *Setx^+/+^ and Setx^-/-^* knockout mice (28 days old) and the results compared with the list of genomic gains and losses (Figure 3C, Supplementary Figure S3B-D, and Supplementary Table S3A-F). Consistent with the AOA2 patient data, the CNCs mapped to gene-rich regions (Figure 3D and E). Strikingly, both in MEFs and cerebellum, the gain regions were significantly enriched for Ensembl genes (*P*=6.5 × 10^−11^ and *P*=1.3 × 10^−02^, respectively, by permutation test). In MEFs, but not in testis and cerebellum, we identified that the gain regions were significantly enriched for ERFS (*P*=8.4 × 10^−11^ by permutation test), which is in agreement with results from the AOA2-P1 fibroblasts. In cerebellum, we found 27% of the loss regions were associated with CFSs (49), while none of the gains/losses from MEFs overlapped with CFSs (Figure 3C). In addition, some of the mouse cerebellum CNCs overlapped with recurrent DNA break clusters (RDCs) identified recently in long neural genes from neural stem/progenitor cells (Figure 3C) (50, 51). Collectively, these results show that loss or inactivation of SETX gives rise to genome-wide chromosomal fragility associated with the transcribed regions of the mammalian genome.

### Differential gene expression in SETX-deficient human and mouse cells

Because CNCs can affect the cellular transcriptome (44) and AOA2 cells exhibit altered gene expression profiles (52), gene expression microarrays of the AOA2 (P1 - P4) and CTRL (C1 - C4) LCLs were analyzed. Expression of a significant number of genes (ranging from 365 to 675 genes) showed a >2-fold change (p<0.05) in the AOA2 LCLs. Comparison of AOA2 LCLs with their controls revealed a total of 832 upregulated and 656 downregulated genes in AOA2 (Supplementary Table S4A and B). Strikingly, the expression of 310 genes (209 and 101 genes were upregulated and downregulated, respectively) was significantly altered in all four AOA2 cell lines. Among these, 27 upregulated and 12 downregulated genes correlated with genomic regions that scored as gains and losses, respectively.

To identify the genes most affected by *SETX* mutations, we carried out RNA-seq with AOA2 (P1-P4) and CTRL (C1-C4) LCLs and found numerous genes that were differentially expressed (upregulated and downregulated) in AOA2 cells (Supplementary Figure S4A-D, and Supplementary Table S5A and B). Comparison of the AOA2 LCLs revealed a core set of 96 genes (55 upregulated and 41 downregulated; >2-fold change) that exhibited the greatest expression differences relative to the controls (Supplementary Figure S4E). Furthermore, gene set enrichment analysis (GSEA) using the ranked test-statistics from the differential expression analysis of CTRL and AOA2 LCLs revealed that biological process terms such as regulation of transcription from RNAPII promoter in response to stress, RNA splicing, regulation of RNA stability and RNA 3’-end processing, were significantly (FDR<0.05) enriched (Supplementary Table S6).

Differential gene expression profiles were also confirmed with *Setx^-/-^* knockout mouse MEFs (684 upregulated genes and 470 downregulated genes) by RNA-seq analyses (Figure 4F, and Supplementary Table S5C and D). Importantly, overlap analyses revealed that the gain regions identified in *Setx^-/-^* MEFs were enriched for upregulated genes (*P*=1.2 × 10^−08^ by permutation test), while loss regions did not show significant overlap with downregulated genes. Taken together, these data indicate that CNCs can contribute to the gene expression changes identified in the absence of SETX.

**Figure 4.**
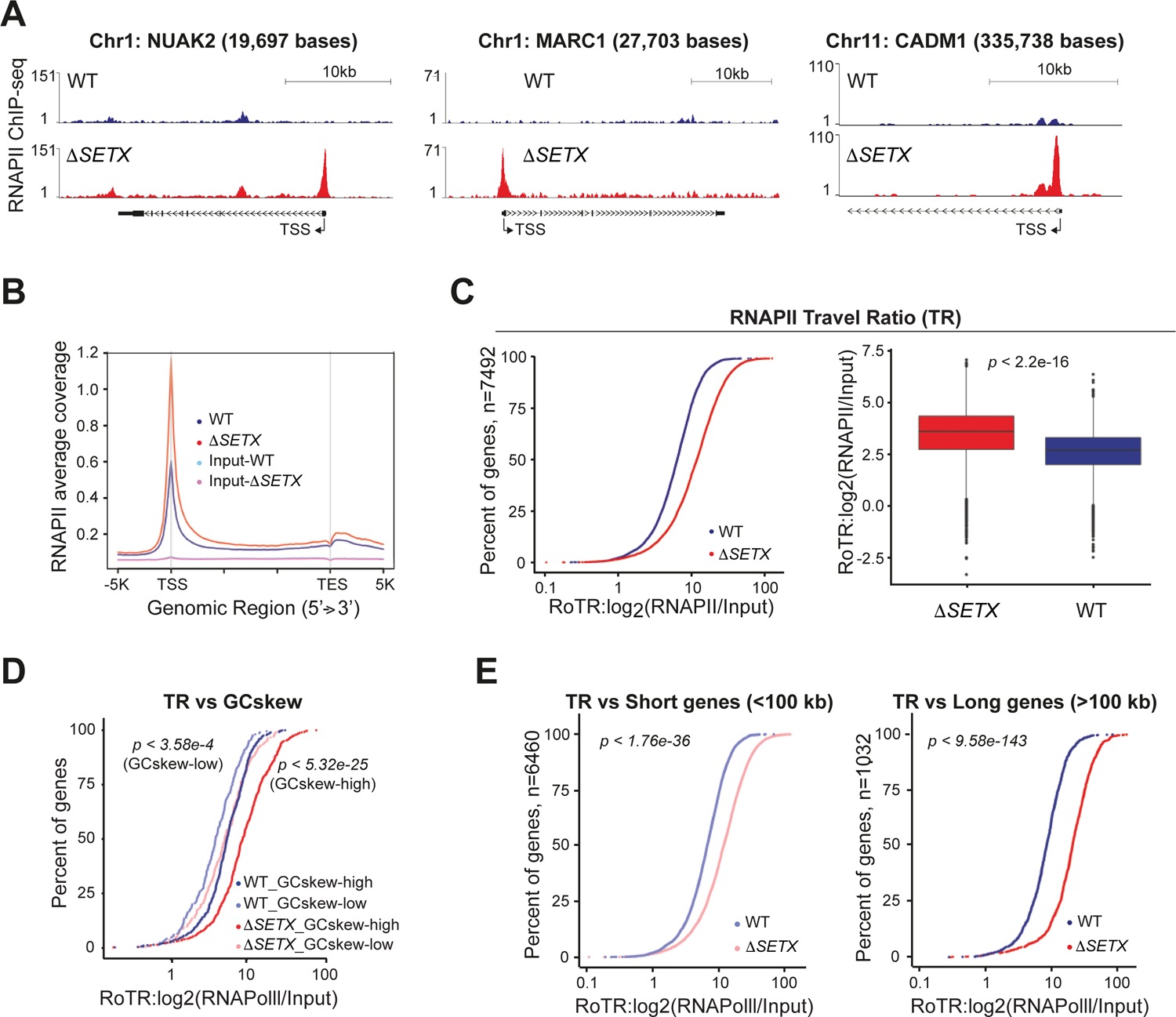
Loss of SETX causes transcription stress. (A) ChIP-seq analyses using RNA polymerase II antibodies were carried out with cross-linked chromatin from HAP1 wild-type (WT) and Δ*SETX* cell lines. Representative genome browser shots of RNAPII binding over the *NUAK2*, *MARC1* and *CADM1* genes are shown. TSS, transcription start site. Scale bars, 10kb. (B) Average RNAPII ChIP-seq coverage profiles for WT and Δ*SETX* are shown, along with input controls. TSS and TES, transcription start and end sites. (C) Left: Cumulative curve of RNAPII ratio of traveling ratios (RoTR; x-axis) for 7492 genes in WT and Δ*SETX*. The y-axis indicates percent of all genes. Right: Box plots show the distribution of RoTR of 7492 genes in WT and Δ*SETX*. P values were calculated using non-parametric Wilcoxon rank sum test. (D) Cumulative curve of RoTR in WT and Δ*SETX* with GCskew-high (n=505) and GCskew-low (n=215) groups that were called based on a 10% and 90% quantile from the GCskew frequency distribution of the entire genic intervals (n=26934). P values were calculated as in (C). (E) Left: Cumulative curve of RoTR of 6460 short genes (<100 kb) in WT and Δ*SETX.* Right: Cumulative curve of RoTR of 1032 long genes (>100 kb) in WT and Δ*SETX*. P values were calculated as in (C).

### Loss of SETX alters the profile of RNAPII across the genome

To investigate the role of SETX in transcription, RNAPII ChIP-sequencing (ChIP-seq) was carried out with human HAP1 wild type (WT) and *SETX* knockout (Δ*SETX*) cells. Strikingly, a genome-wide increase in RNAPII levels over transcription start sites (TSS) was observed in the Δ*SETX* cells (Figure 4A and B). To quantitatively analyze how SETX affects transcription at the genomic level, we measured the traveling ratio (TR) that compares the ratio between RNAPII density in the promoter-proximal region (−30 to +300 bp relative to the TSS) and the remaining length of the gene (53). The TR was found to be globally higher in Δ*SETX* cells compared to WT and input controls (Fig S5A and B). When we focused our analysis on 7,492 genes (Supplementary Table S7) with clear RNAPII peaks at the promoter-proximal region (active genes), we observed that these genes exhibited a higher TR in Δ*SETX* compared to WT (Figure 4C and Supplementary Figure S5C). Collectively, these data show that SETX inactivation affects RNAPII progression throughout the genome.

Base composition analyses revealed significantly higher GC-richness and GCskew (strand asymmetry in the distribution of guanines and cytosines) at the promoter-proximal regions of these 7,492 genes (Supplementary Figure S5D). To investigate how GCskew affects TR, we calculated GCskew of genic regions (n = 26934), and subsequently “GCskew-high” and “GCskew-low” groups were called based on a 10% and 90% quantile from the distribution of the GCskew values. Strikingly, the differences in TR between Δ*SETX* and WT was significantly higher for “GCskew-high” genes (n = 505) compared to “GCskew-low” (n = 215) genes (Figure 4D).

Previously, it was shown that transcription through regions of GCskew leads to the formation of long and stable R-loops (8). Consistent with this, DRIP analyses revealed the accumulation of R-loops at six different genes (*NUAK2*, *CADM1*, *TRIM33*, *SOD1*, *MARC1* and *DDIT4L*) identified to have transcription stress in Δ*SETX* (Supplementary Figure S5E). Importantly, RNaseH treatment, but not RNaseA treatment, abolished R-loop accumulation. The negative control gene, *SNRPN,* did not exhibit R-loop accumulation.

Because long genes (CFSs) and highly transcribed short genes (e.g. histone genes) are prone to transcription stress and R-loop formation (45,54,55), we next focused on the relationship between gene length and TR of the 7,492 genes. We classified 6,460 and 1,032 genes as short (<100 kb) and long (>100 kb) genes, respectively. We found that the TR was significantly higher for both long and short genes in Δ*SETX* compared to WT (Figure 4E), although the TR difference was greater in long compared with short genes. Similar results were observed when the genes were stratified into short (shortest 20%), medium (middle 40-60%) and long (longest 20%) groupings based on quantiles from the gene-width distributions. The results confirmed that the TR differences were much higher in long genes (Supplementary Figure S5F).

In agreement with the findings from AOA2 and *Setx^-/-^* MEFs, differential gene expression profiles were also observed in human HAP1 Δ*SETX* cells (248 upregulated genes and 364 downregulated genes) by RNA-seq analyses (Supplementary Figure S4G, and Supplementary Table S5E and F).

Together, these data lead us to conclude that SETX-defective cells exhibit transcription stress (altered RNAPII transcription) that drives R-loop accumulation and affects gene expression.

### Transcription stress correlates with chromosome instability upon SETX deficiency

To confirm and extend the results observed with AOA2 LCLs, we measured CNCs in the Δ*SETX* cell line and identified 17 and 14 genomic abnormalities of which 13 (9 losses and 4 gains) were common to the two independent experiments (Supplementary Figure S6A and Supplementary Table S8). Consistent with them arising from chromosome instability, 11 of the 13 CNCs mapped to genes and 6 coincided with CFSs. Genomic abnormalities observed in Chromosome 16 in Δ*SETX* encompassing the FRA16D locus (WWOX) is shown in Figure 5A. Others were generally located in gene-rich regions and showed increased RNAPII pausing at gene promoters. The genomic gain at chromosome 17 is shown as an example (Figure 6B).

**Figure 5.**
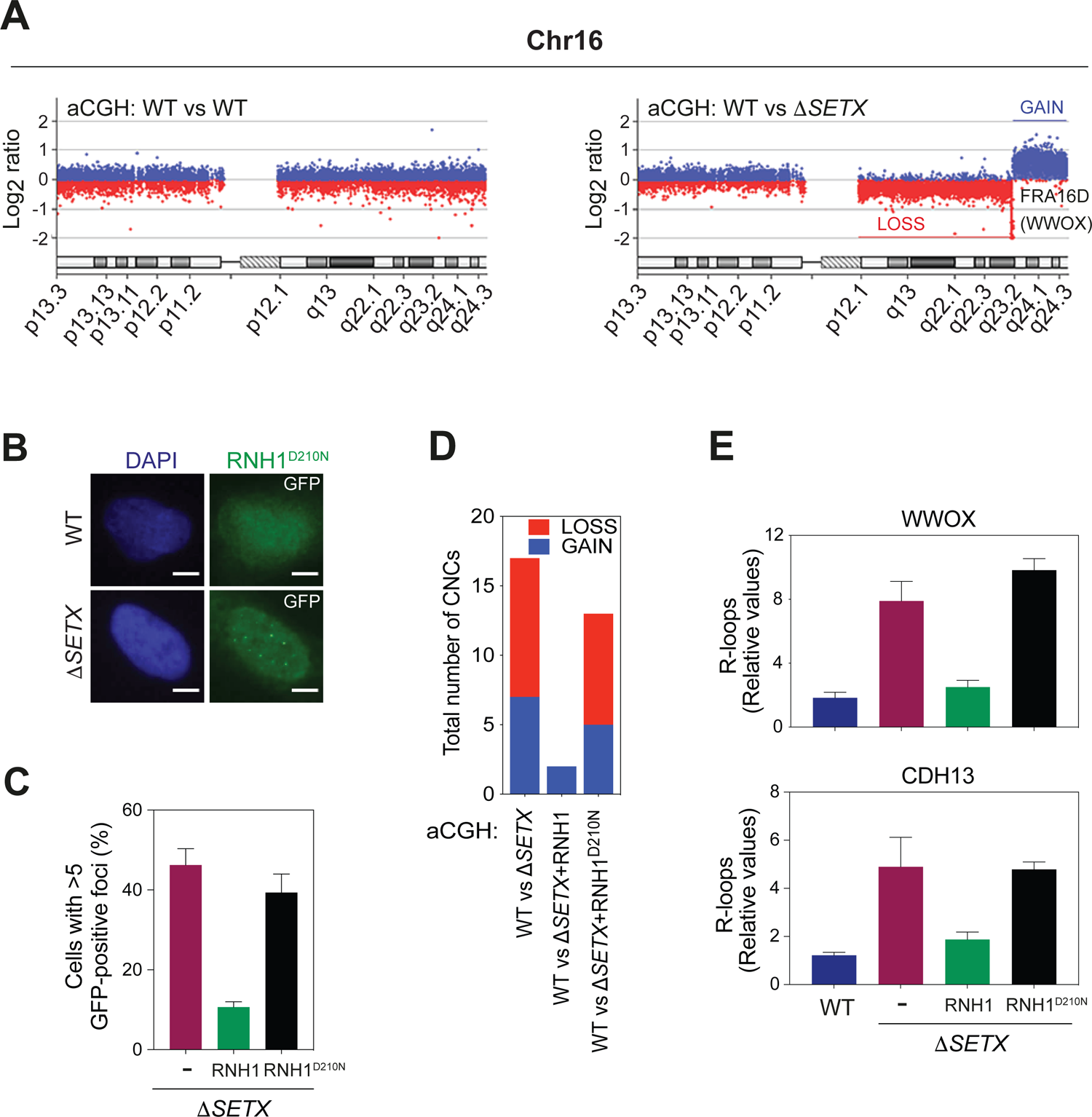
SETX suppresses transcription stress that promotes chromosome instability. (A) aCGH profile of chromosome 16 (Chr16) in WT versus WT condition (left) and WT versus Δ*SETX* (right). The WWOX/FRA16D locus at q23.1 is lost. Other gain and loss regions are marked in blue and red lines, respectively. (B) Detection of R-loops in WT and Δ*SETX* cells by immunofluorescence using catalytically-inactive RNaseH1 (RNH1^D210N^) fused to GFP. Nuclear DNA was stained with DAPI (blue). Representative images are shown. GFP, green fluorescent protein. Scale bar, 5 μm. (C) Quantification of GFP-positive RNH1 foci in Δ*SETX* cells transfected with RNH1 or RNH1^D210N^. Data represent the mean ± s.e.m of three independent experiments with >100 cells per condition. (D) aCGH analyses were carried out with WT vs Δ*SETX*, WT vs Δ*SETX* + RNH1 and WT vs Δ*SETX* + RNH1^D210N^. The graph depicts the number of CNCs identified in each experiment. Chromosome regions with gain (blue) and loss (red), are indicated. Data represent the mean of two independent experiments. (E) DRIP-qPCR analysis using genomic DNA from WT, Δ*SETX*, Δ*SETX* + RNH1 and Δ*SETX* + RNH1^D210N^. R-loop regions at *WWOX* (top) and *CDH13* (bottom) were analysed. Data represent the mean ± s.e.m of three independent experiments.

**Figure 6.**
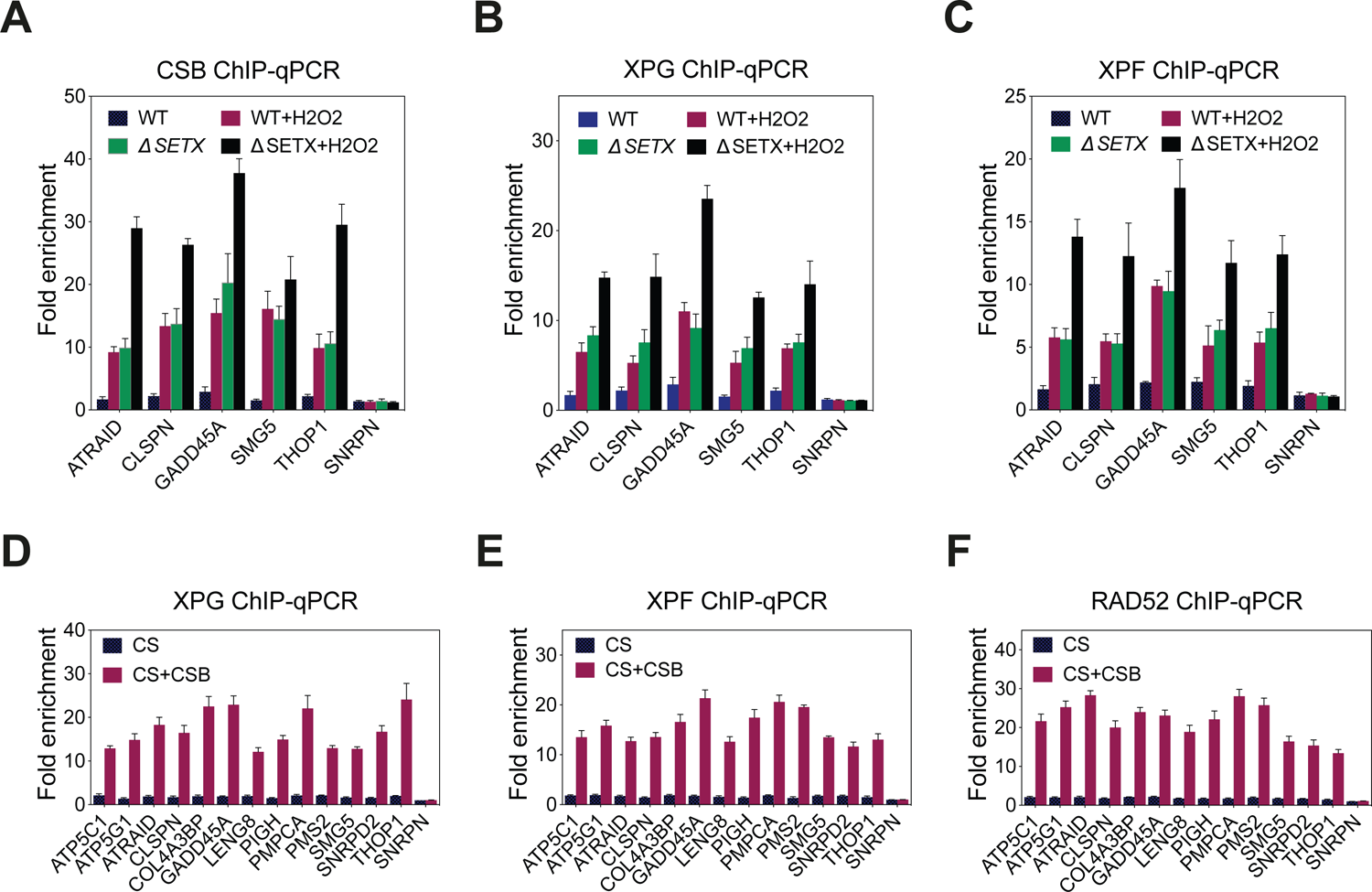
CSB promotes error-prone repair of TSS-associated R-loops in the absence of SETX. (A) ChIP assays were carried out using cross-linked chromatin from HAP1 WT and Δ*SETX* cells treated with or without H_2_O_2_. Rabbit polyclonal antibody against CSB was used for ChIP. TSS regions from the indicated genes were analysed by qPCR. Data represent the mean ± s.e.m of three independent experiments. (B) As (A), using XPG monoclonal antibody. (C) As (A), using XPF rabbit polyclonal antibody. (D) As (B), except cross-linked chromatin from CS1ANsv (CS) and CS cells stably expressing GFP tagged CSB from a BAC (CS + CSB) were used for ChIP. (E) As (D), using XPF rabbit polyclonal antibody. (F) As (D), using RAD52 rabbit polyclonal antibody.

SETX-related genomic instability was associated with the formation of R-loops as determined by immunofluorescent analysis in Δ*SETX* cells overexpressing a catalytic-inactive mutant of GFP-tagged RNaseH1 (RNH1^D210N^) (Figure 5B and C). Consistent with this, 13 genomic alterations (8 losses and 5 gains) were identified in two independent aCGH experiments carried out with WT vs Δ*SETX* + RNH1^D210N^ cells (Supplementary Figure S6A). Eleven of these were similar to aberrations identified in the earlier aCGH experiments with WT vs Δ*SETX* (Figure 5D). On the other hand, aCGH experiments with WT vs Δ*SETX* + RNH1 revealed only 2 non-specific (both in Y chromosome) genomic aberrations. DRIP analysis of the *WWOX* and *CDH13* fragile site loci confirmed the accumulation of R-loops in Δ*SETX and* Δ*SETX*+RNH1^D210N^ cells, but not in Δ*SETX* + RNH1 cells (Figure 5E).

Furthermore, these CFS loci accumulated significant levels of DNA damage, as measured by γ-H2AX ChIP analysis (Supplementary Figure S6C). Together, these results confirm that transcription stress causes chromosome instability in cells lacking SETX.

### Transcription coupled repair promotes genomic instability

Unscheduled or unwanted R-loops that form co-transcriptionally are prone to DNA breakage and genomic rearrangements (56, 57). Transcription-coupled repair (TCR) proteins were shown to be involved in loop cleavage, which ultimately leads to DNA double-strand break formation and drives genome instability (21). We therefore investigated whether TCR proteins are recruited to the TSS of RNAPII transcribed genes that showed elevated RNAPII pausing and increased R-loop formation upon SETX deficiency. In five genes tested, ChIP analysis revealed an enrichment of the TCR proteins CSB, XPG and XPF at the TSS in Δ*SETX* compared to WT cells (Figure 6A-C). Treatment of the cells with H_2_O_2_ further increased the occupancy of TCR proteins at these sites (Figure 6A-C), consistent with the observation that oxidative stress stabilizes promoter-proximal paused RNAPII (58). Using Cockayne Syndrome (CS) and CSB-complemented cells (CS+CSB), we found that the absence of CSB resulted in a failure to recruit XPG and XPF endonucleases for the processing of R-loops at TSS of genes containing paused RNAPII (Figure 6D and E).

The homologous recombination protein RAD52 promotes DNA repair in actively transcribed regions by transcription-coupled homologous recombination (59, 60). Consistent with this, ChIP analyses revealed that the absence of CSB led to a failure to recruit RAD52 to these TSSs (Figure 6F).

To further understand the relationships between transcription coupled repair and SETX in R-loop processing, we determined the levels of DNA damage at the TSS regions of genes with paused RNAPII. ChIP analyses revealed an enrichment of γH2AX at the TSS of these genes in SETX-depleted MRC5 cells compared to controls (Figure 7A). We also detected increased levels of γH2AX at the TSS of these genes in CS, but not CS complemented (CS+CSB) cells. The levels of γH2AX were consistently reduced in CS cells compared with SETX-depleted cells, and depletion of SETX from the CS cells led to a further increase in γH2AX levels (Figure 7A). Importantly, the levels of γH2AX accumulation detected in control or SETX-depleted MRC5, CS and CS+CSB cells correlated with the levels of R-loops at the TSS of these genes (Figure 7B). These results indicate that the TCR pathway and SETX act independently to remove R-loops at TSS of RNAPII transcribed genes. Consistent with this, we found that SETX-depleted CS cells exhibit high frequencies of bulky DNA anaphase bridges and lagging chromosomes (Figure 7C and D). Quantification revealed that the occurrence of these chromosome aberrations was further increased upon H_2_O_2_ treatment that stabilizes promoter-proximal paused RNAPII (Figure 7D).

**Figure 7.**
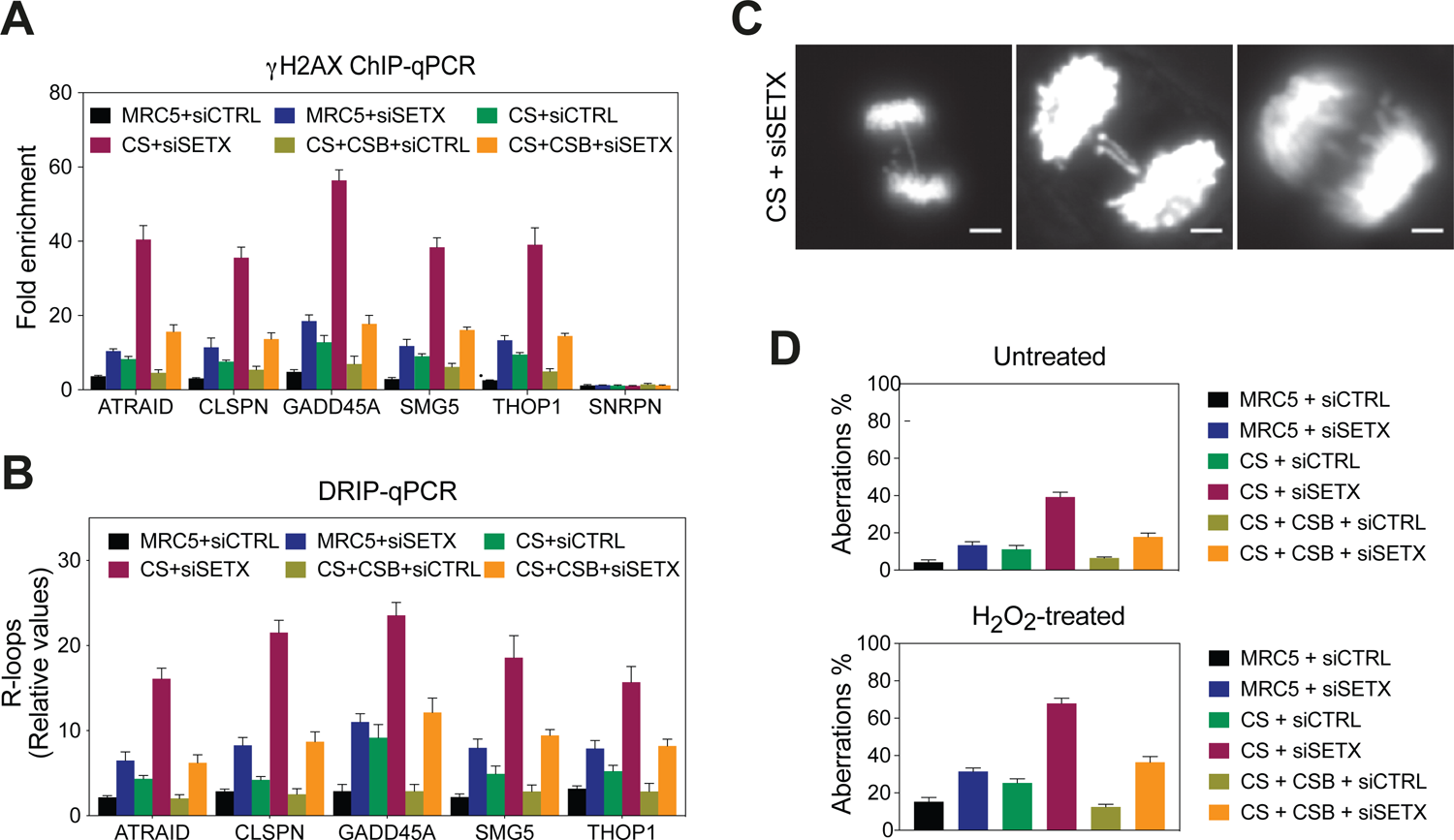
CSB induced DNA damage in SETX-deficient cells. (A) ChIP assays were carried out using cross-linked chromatin from the indicated cells treated with control siRNA (siCTRL) or SETX siRNA (siSETX). TSS regions from the indicated genes were analysed by q-PCR. Data represent the mean ± s.e.m of three independent experiments. (B) DRIP assays were carried out using cross-linked chromatin from the indicated cells treated with siCTRL or siSETX. TSS regions from the indicated genes were analysed by q-PCR. Data represent the mean ± s.e.m of three independent experiments. (C) Representative images of chromosome aberrations (bulky DNA bridges and lagging chromosomes) in SETX-depleted CS cells. DNA was stained with DAPI. Scale bar, 5 μm. (D) Quantification of the percentage of chromosome aberrations in the indicated cells treated with or without H_2_O_2_. 300 mitotic cells were analysed. Data represent the mean ± s.e.m of three independent experiments.

## DISCUSSION

Our results demonstrate that the loss of SETX causes transcription stress, increased R-loop formation and chromosomal instability at RNAPII transcribed genes and fragile sites. SETX promotes R-loop repair (RLR) to remove R-loops that form near the promoters of RNAPII transcribed genes, and thereby controls transcription and maintains genome stability (Supplementary Figure S7). In the absence of SETX aberrant R-loops are incised by transcription coupled repair nucleases, and acted upon by recombination proteins, resulting in error-prone repair and genomic instability (Supplementary Figure S7).

Active gene promoters are known to represent major hotspots for R-loop formation, as ∼60% of R-loops map to promoter-proximal regions (6-8,16). We have shown that SETX acts at the promoter-proximal regions of RNAPII transcribed genes to control transcription stress-induced R-loop accumulation and prevent genome-wide chromosomal instability. Our results are consistent with previous observations showing that promoters containing R-loops that arise due to SETX-deficiency are resistant to cytosine methylation by DNA methyltransferase 1 (DNMT1) and exhibit increased transcription (32). The underlying basis for this lack of methylation is that DNMT1 preferentially binds double-stranded DNA, but not RNA-DNA hybrids (32). Interestingly, SETX-mediated R-loop removal is important for chromosome stability, in particular at common fragile sites, long genes and highly transcribed genes, all of which are prone to replication/transcription stress. The mis-regulation of vital transcripts, in addition to the observed chromosome instability, is may contribute to disease progression in AOA2.

The fate of paused RNAPII remains enigmatic. Early observations led to the proposal that RNAPII backtracking at pause sites provides a free RNA 3’ end for the RNA exosome, resulting in degradation of the RNA transcript and transcription termination (61). However, increased RNAPII pausing and backtracking also leads to R-loop formation and genome instability (62, 63). Transcription problems arising due to backtracking are resolved by TFIIS (also known as TCEA1) and intrinsic transcript cleavage by RNAPII. Using TFIIS mutant cells, it was recently shown that R-loops form at the anterior side of backtracked RNAPII and trigger genome instability (63). Possible interactions between TFIIS and SETX could provide a link between RNAPII backtracking and the R-loop resolution machinery, supporting the notion that SETX plays an important role in the removal of R-loops formed at the anterior side of RNAPII.

In yeast and mammals, Sen1/SETX facilitates pause-dependent transcriptional termination at specific RNAPII transcribed genes (9,19,64–66). Transcription termination requires a functional polyadenylation signal (PAS) and either downstream pause sites or co-transcriptional cleavage sequences together with 3’ transcript degradation by the 5’-3’ exonuclease Rat1 (in budding yeast) and XRN2 (in humans) (67–71). Interestingly, the terminator regions of PAS-dependent genes, which are devoid of GC skewness but contain G-rich pause sites, facilitate RNAPII pausing and contribute to R-loop formation (16). These R-loop structures are resolved by SETX and the nascent RNA is degraded by XRN2 leading to efficient transcription termination (9).

Recent genome-wide analyses revealed that the majority of R-loops map to promotor proximal regions with less than 7% of R-loops found at terminator regions (6). In the present work, we observed only a slight increase in RNAPII changes at the terminator-proximal regions upon SETX deficiency. These results are consistent with the existence of several SETX-independent transcription termination mechanisms in human cells. For example, it has been shown that the SCAF4 (SR-related CTD associated factor 4) and SCAF8 proteins are required for efficient RNAPII transcription termination (72). The absence of SCAF4/SCAF8 affects premature PAS site selection and termination in more than 1300 genes. Notably, these proteins were found to interact with 3’ end-processing factors, including members of the cleavage and polyadenylation specificity factor (CPSF) complex, but not with SETX or the RNA exosome. R-loop associated genomic instability has been reported in CPSF and other 3’ end-processing mutants, suggesting links between R-loops, 3’ end-processing and transcription termination (20,73,74). Recent observations also suggest that R-loops induce low levels of antisense transcription at terminator regions of short and ubiquitously expressed genes, resulting in the formation of transient double-stranded RNA and triggering an RNAi response to induce the formation of repressive chromatin marks that enhance RNAPII pausing and transcription termination (4). Moreover, increased formation of insulator regions/gene loops over R-loop containing gene terminators by CTCF-cohesin complex contributes to efficient transcription termination (16, 75).

Genome instability in the adult brain leads to impaired neural development and neurodegeneration (76, 77). The central nervous system is particularly vulnerable to oxidative stress due to the high levels of oxygen consumption, low levels of antioxidant enzymes and the terminally differentiated state of neurons. Indeed, oxidative stress represents a major cause of neuropathology underlying a variety of neurodegenerative diseases. Moreover, DNA breaks are associated with cognitive impairment in neurodegenerative conditions, such as Alzheimer’s disease and ALS. Consequently, mutations in many DNA single- and double-strand break repair proteins have been linked to multiple neurodegenerative diseases including other subtypes of AOA. Recent studies indicate that oxidative stress (i.e. treatment with a low dose of H_2_O_2_) causes a rapid and genome-wide increase (up to 5-fold) in promoter-proximal paused RNAPII (58). The paused RNAPII begin to creep into gene bodies after ∼10 min, and this was correlated with the loss of negative elongation factor (NELF) at promoters. Importantly, the effects caused by peroxide treatment were not dependent on DNA damage. These observations are consistent with the results presented here and provide an explanation for the hypersensitivity of AOA2 cells to H_2_O_2_ treatment, together with altered gene expression profiles and genomic instability, which occurs as a result of transcription stress due to SETX-deficiency.

In conclusion, our findings show that AOA2 is a transcription stress-related disorder and that SETX is necessary to preserve the integrity of transcription. The absence of SETX leads to R-loop accumulation, and these serve as sites of incision by the TCR proteins XPG and XPF leading to error-prone repair and DNA damage. Targeting of these endonucleases is dependent upon CSB. Other neurological diseases, including Tremor-Ataxia Syndrome (TAS), Autosomal Dominant proximal Spinal Muscular Atrophy (ADSMA) and Charcot-Marie-Tooth (CMT) disease have also been linked with mutations in the *SETX* gene (78–80). Further studies will provide invaluable insights that will allow us to piece together the relationships between transcription stress and human disease.

## ACKNOWLEDGEMENTS

We thank members of the West lab for their help and encouragement, Jesper Svejstrup, Jiri Lukas, Maria Teresa Bassi and Luciana Chessa for the cell lines, Stephen H. Leppla for the S9.6 antibody, the CRUK Manchester Institute Microarray Service for expression array analysis and the Francis Crick Institute’s Cell Services, Flow Cytometry and Advanced Sequencing facilities for cell culture, Imagestream and sequencing support. This work was supported by the Francis Crick Institute (FC10212), the European Research Council (ERC-ADG-249145 and ERC-ADG-666400) and the Louis-Jeantet Foundation. The Francis Crick Institute receives core funding from Cancer Research UK, the Medical Research Council, and the Wellcome Trust. R.K was the recipient of fellowships from the Swiss National Science Foundation (PBZHP3-138777 and PA00P3-142187).

## AUTHOR CONTRIBUTIONS

Conceptualization, R.K. and S.C.W.; Methodology, R.K.; Formal Analysis, R.K., R.M., T.K., M.M.E. and P.C.; Investigation, R.K., T.K., A.B., B.F., F.Y.; Resources, O.B., M.F.L.; Writing, R.K. and S.C.W.; Funding Acquisition, S.C.W.; Supervision, S.C.W., A.S., and A.K.

## Supplementary Figure Legends

## METHODS

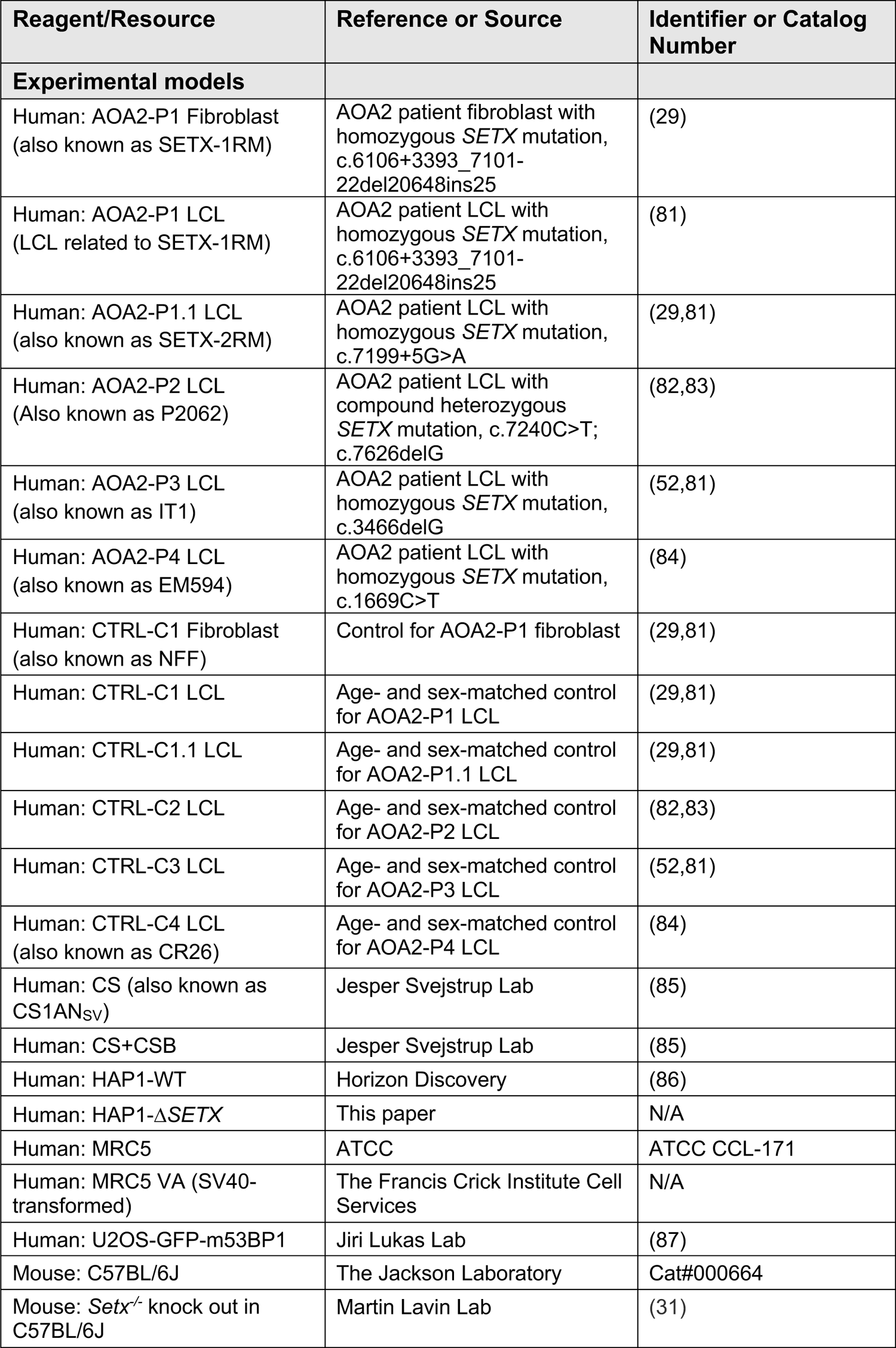
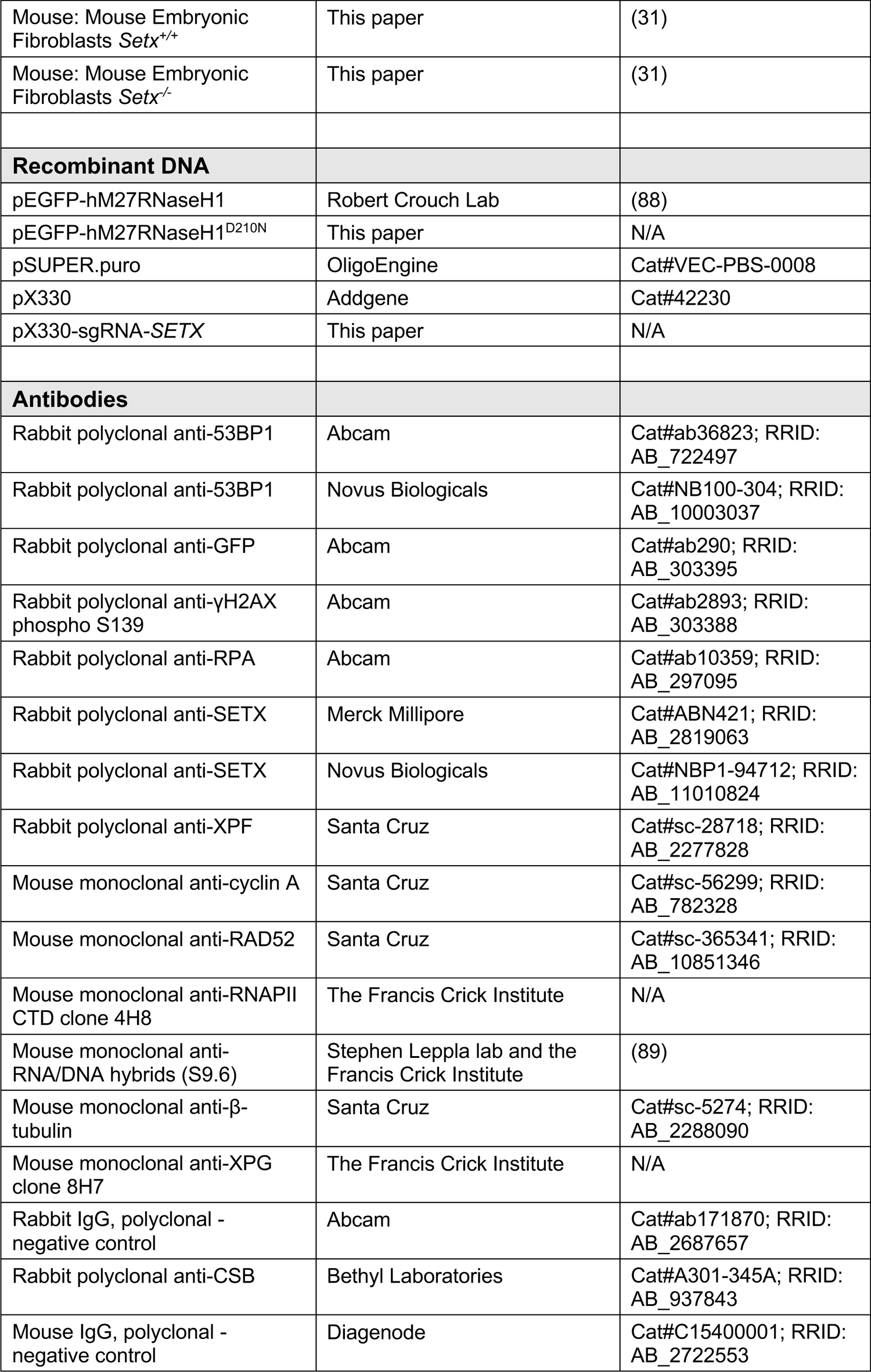
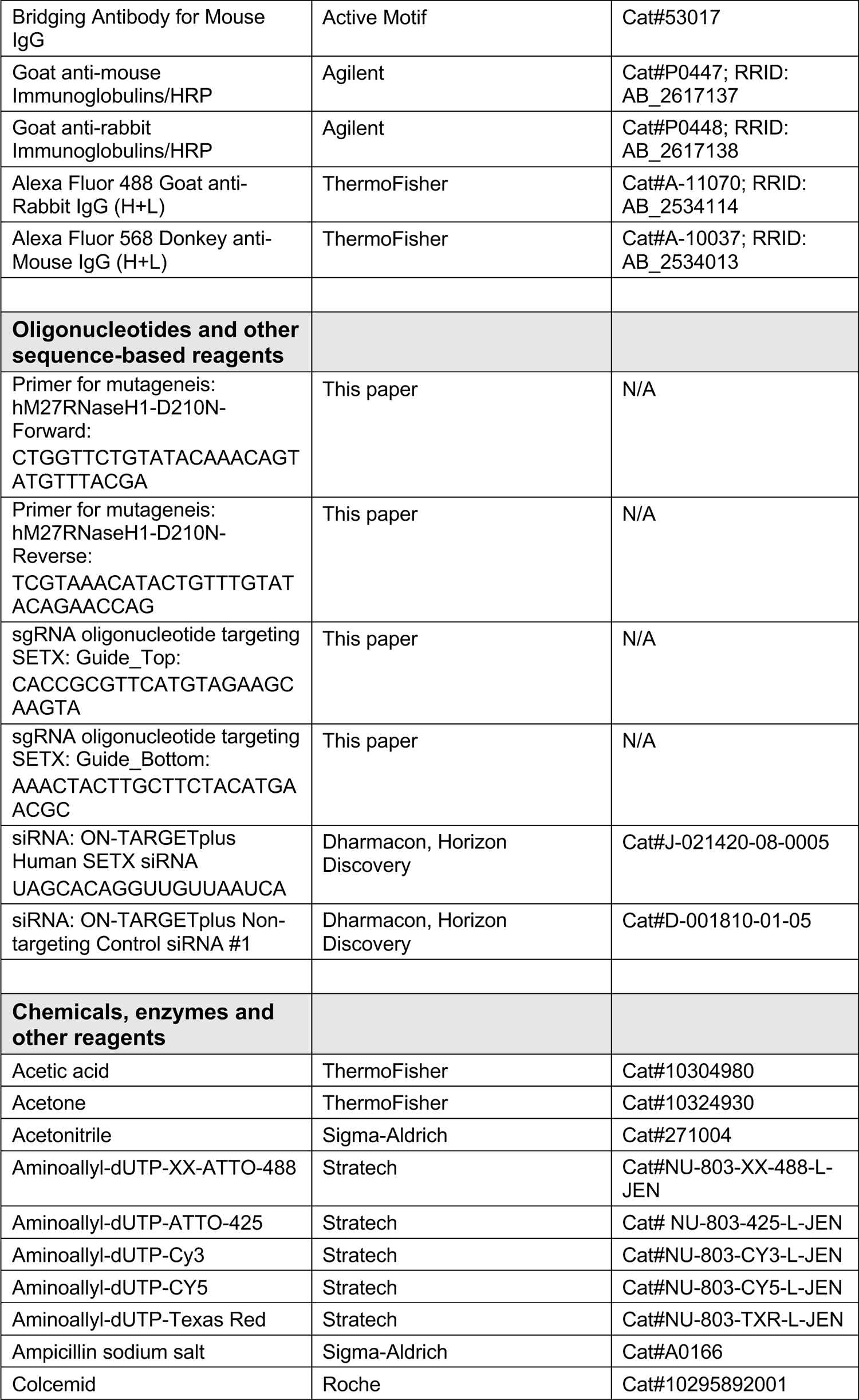
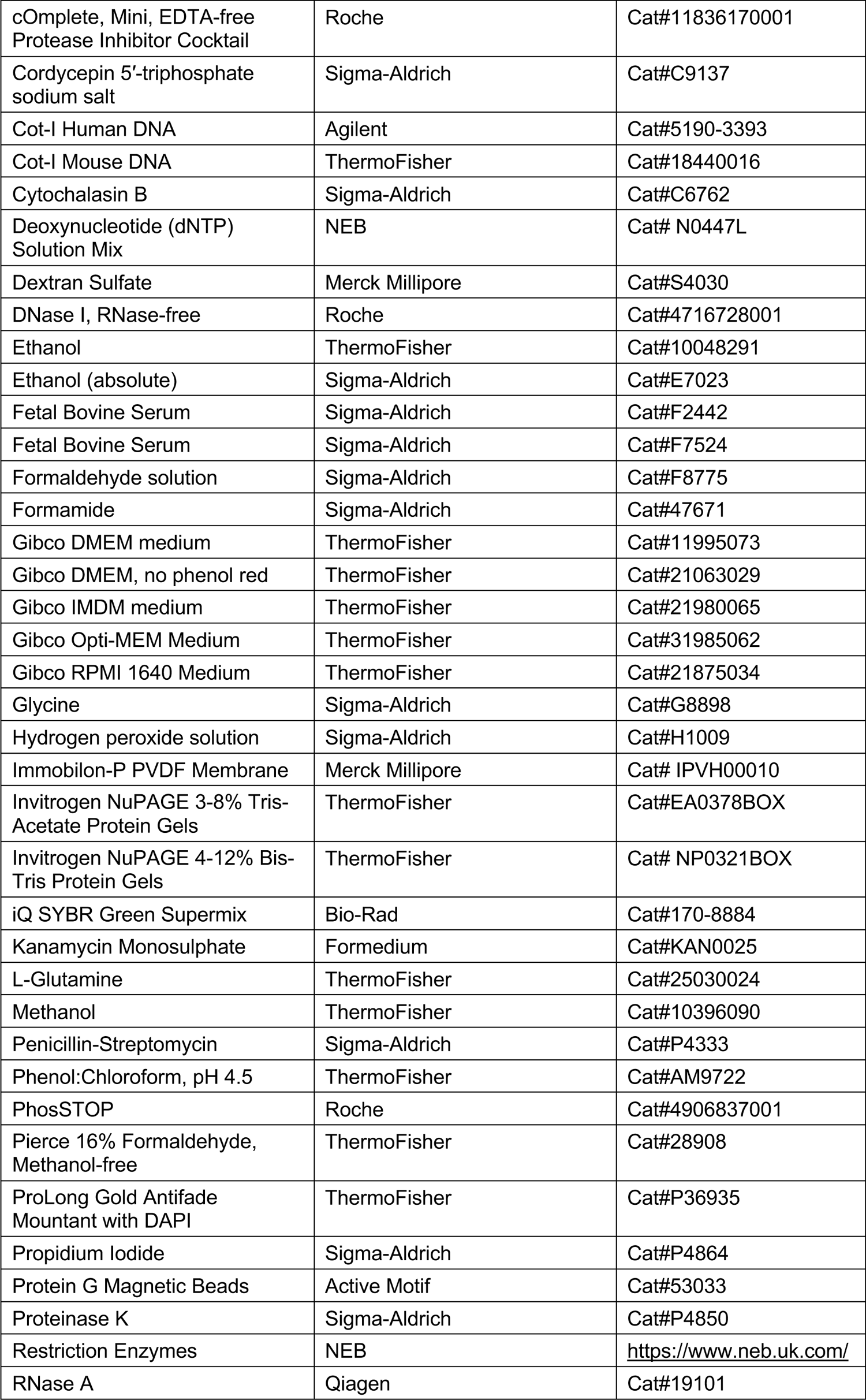
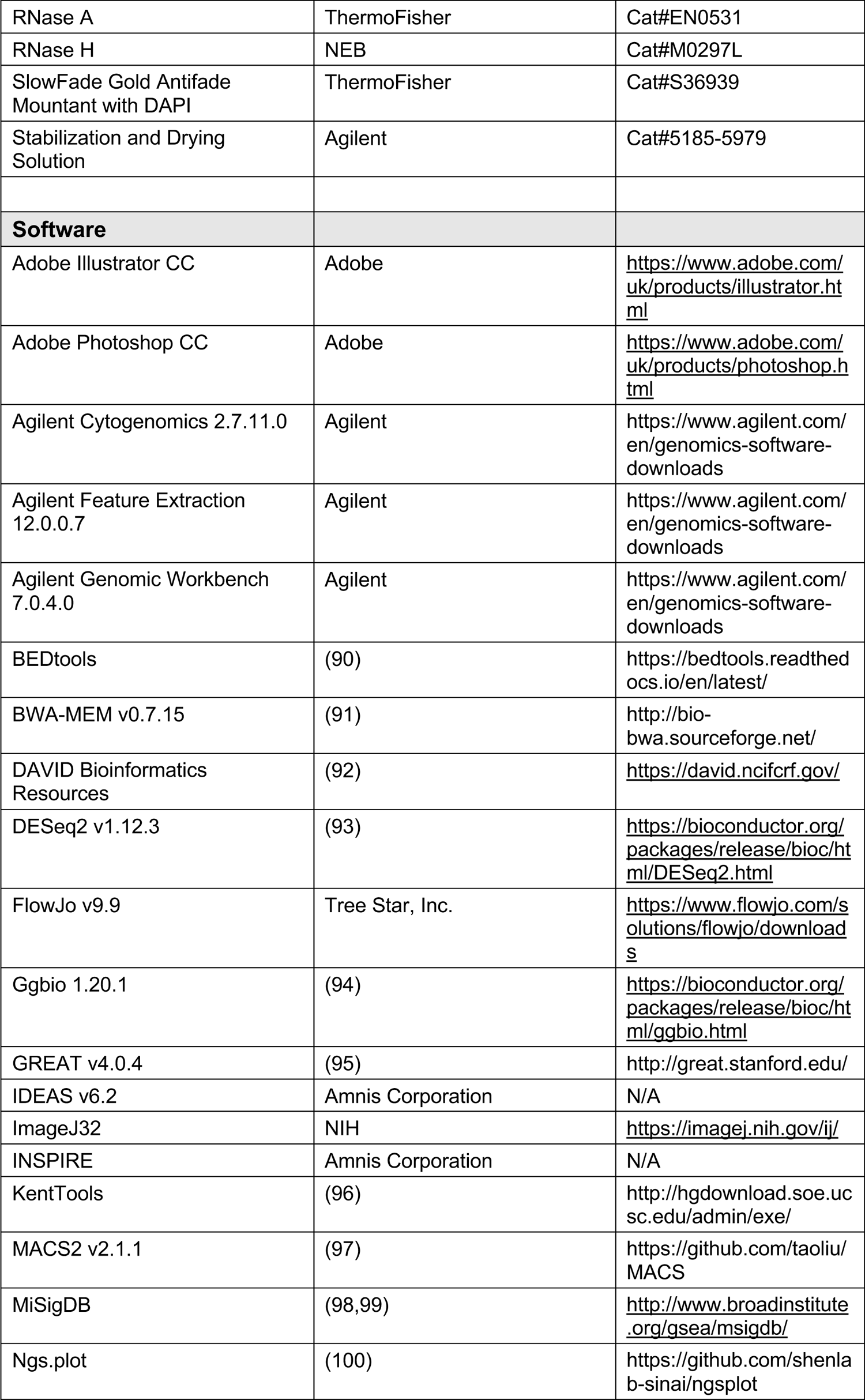
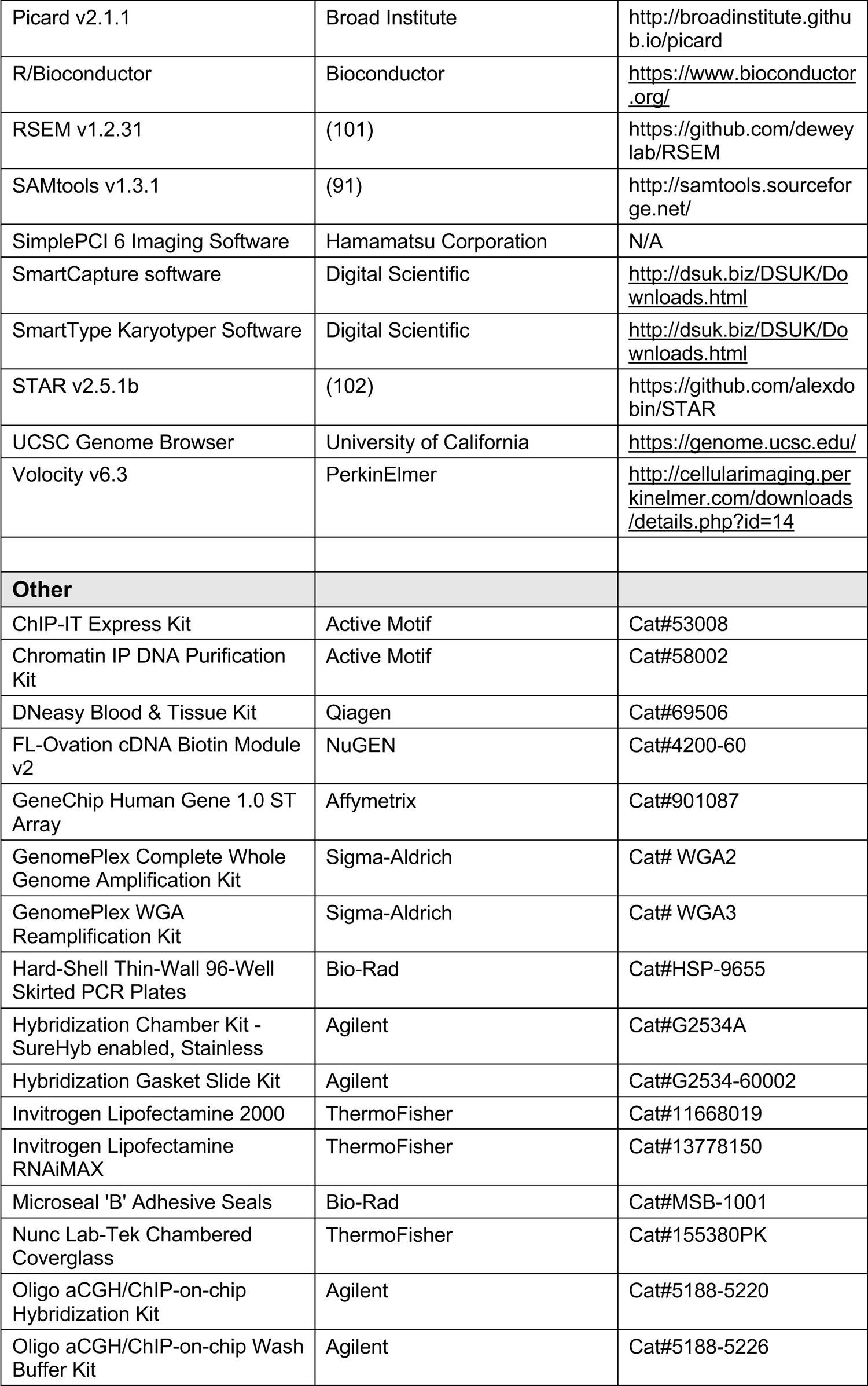
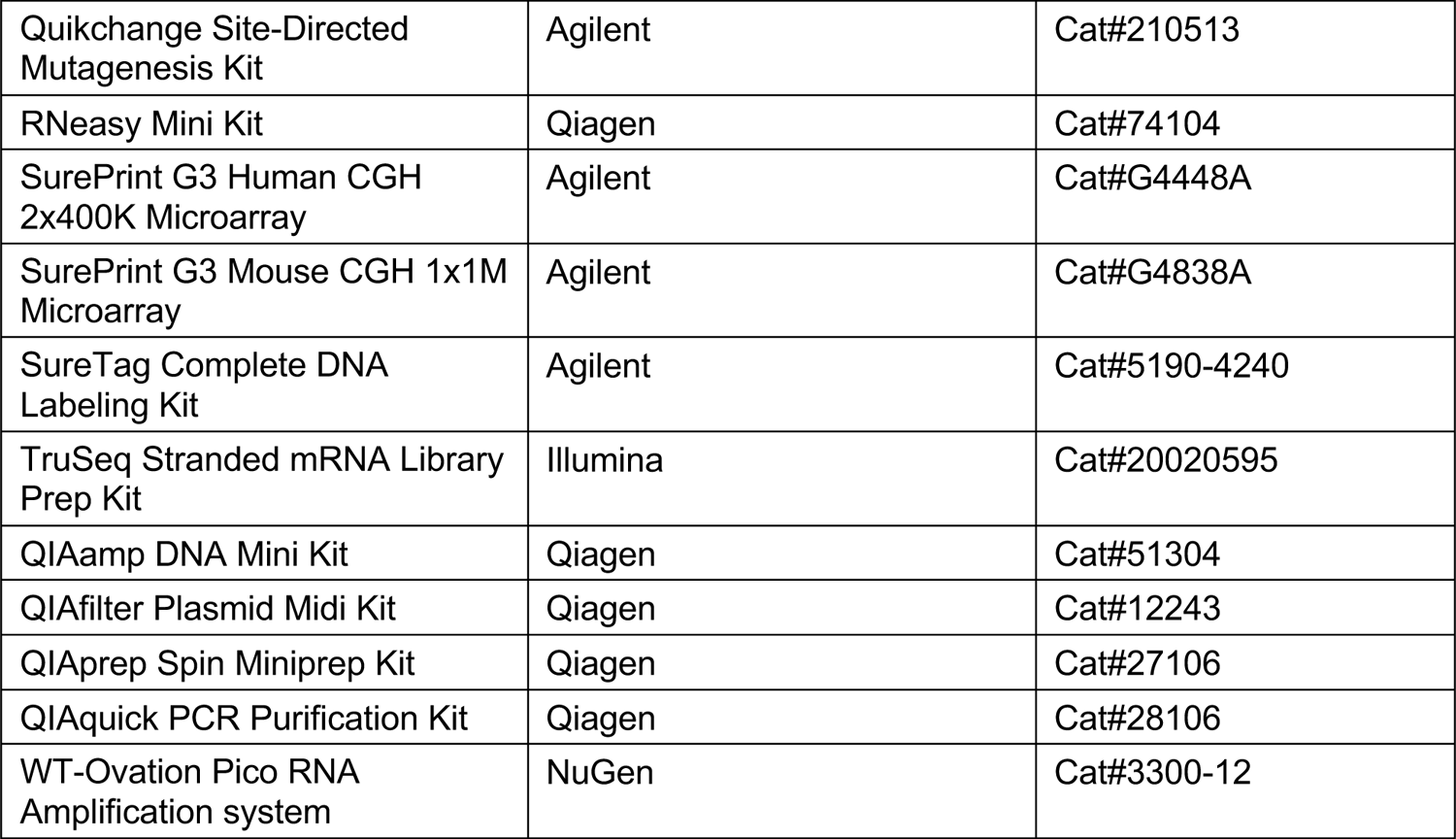

### Method Detail Cell culture

MRC5 (MRC5VA), CS (CS1AN_SV_; Cockayne syndrome B [CSB] fibroblast SV40 immortalized) and CS + CSB (CS cells stably expressing GFP tagged CSB from a BAC) cells were cultured in DMEM (ThermoFisher) supplemented with 10% fetal bovine serum (FBS) and 1% penicillin/streptomycin (P/S). U2OS cells stably expressing GFP tagged mouse 53BP1 (U2OS-GFP-m53BP1), were grown in DMEM supplemented with 10% FBS, 1% P/S and 400 μg/ml G418 (ThermoFisher). Human HAP1 WT and Δ*SETX* cells were grown in IMDM medium (ThermoFisher) plus 10% FBS and 1% P/S. Lymphoblastoid cell lines (LCLs) from control and AOA2 patients were cultured in RPMI 1640 medium (ThermoFisher) containing 20% FBS, 1% P/S and 2 mM L-glutamine (ThermoFisher). Human foreskin fibroblasts (HFF; control) and fibroblasts from an AOA2 patient were grown in DMEM supplemented with 15% FBS and 1% P/S. All human cell lines were grown at 37°C and in a humidified atmosphere containing 5% CO_2_. *Setx*^+/+^ and *Setx*^-/-^ mouse embryonic fibroblasts (MEFs) were cultured at 37°C with 5% CO_2_ and 5% O_2_ in DMEM supplemented with 15% FBS (F2442; Sigma) and 1% P/S. Murine tissues were prepared as previously described (31). All animal experiments were approved by the QIMR Berghofer Medical Research Institute Animal Ethics Committee, The University of Queensland, Australia. All cell lines used in this study were confirmed to be mycoplasma-free and were authenticated by STR profiling and PCR based analysis.

### Mouse embryonic fibroblasts isolation and culture

For generation of *Setx*^-/-^ MEFs*, Setx*^+/-^ parent mice were combined for 12 h for copulation. Pregnant females at 13.5 days gestation were subjected to euthanasia under anesthesia. Individual embryos were collected by uterine dissection, rinsed in PBS followed by removal of head, heart and liver. The remaining embryo was minced using sterile scalpel blades and incubated in trypsin at 37°C for 20 min. The tryptic digest was centrifuged and the supernatant discarded. Pelleted cells were suspended in DMEM, 15% FBS and 1% P/S and plated for cell culture. Cells were maintained using a standard 3T3 protocol.

### Plasmids

A short guide RNA (sgRNA) targeting a coding sequence of exon 4 in *SETX* was inserted into the BbsI site of pX330 (plasmid 42230, Addgene) as described (103, 104). The sgRNA oligonucleotides are listed in the reagent and resource table. The plasmid EGFP-N2 containing hM27RNaseH1 (GFP-RNH1) (88), lacking a mitochondrial localisation sequence, was kindly provided by Robert Crouch. The catalytic dead mutant of RNH1, RNH1^D210N^, was generated using the Quikchange Site-Directed Mutagenesis Kit (Agilent).

### Transfection

Transfection of small interfering RNA (siRNA) was carried out using Lipofectamine RNAiMAX according to the manufacturer’s instructions (Invitrogen). For siRNA transfection, cells were seeded at 30% confluence, grown for 24 h, and transfected at a final siRNA concentration of 40 nM. Transfection was repeated 24 h after the first transfection. Unless indicated otherwise, cells were analysed 72 h after the first transfection. For plasmid transfection, cells were seeded at 50% confluence, incubated for 24 h, and transfected with 2 – 4 μg of plasmid DNA using Lipofectamine 2000 based on the manufacturer’s instructions. Unless indicated otherwise, cells were analysed 48 h after transfection.

### Cell line manipulation and generation

To generate *SETX* knockouts in the HAP1 line, cells were transiently transfected with pX330 containing a *SETX*-specific sgRNA sequence, along with pSUPER.puro (Oligoengine) at a 7:1 ratio. After 24 h, cells were selected with 1 μg/ml puromycin (ThermoFisher) for 2 days before the cells were harvested and seeded as single colonies. Initially, clones were selected on the basis of a negative signal for SETX by western blotting. The selected clones were then sequence verified by Sanger sequencing.

### Western blotting

Cells were harvested and resuspended in Iysis buffer (50 mM Tris-HCl pH 7.5, 150 mM NaCl, 0.5% NP-40, 10% glycerol, 1 mM dithiothreitol) supplemented with protease and phosphatase inhibitors (Roche). After a 30 min incubation on ice, the lysates were treated with RNase-free DNase I (10 U, Roche) for 30 min at room temperature (RT), and then clarified by centrifugation at 14,000 rpm for 30 min. Whole cell lysates were subjected to SDS-PAGE and analyzed by Western botting. For western blots the following primary antibodies were used: rabbit anti-SETX (ABN421; 1:2000), rabbit anti-CSB (A301-345A; 1:1000), rabbit anti-GFP (ab290; 1:2000) and mouse anti-β-Tubulin (sc-5274; 1:2000). The secondary antibodies used were goat anti-rabbit Immunoglobulins/HRP and goat anti-mouse Immunoglobulins/HRP (Dako).

### Immunofluorescence

Cells grown on glass coverslips were fixed with PTEMF buffer (20 mM PIPES pH 6.8, 0.2% Triton X-100, 10 mM EGTA, 1 mM MgCl_2_ and 4% formaldehyde [methanol-free]) for 10 min at RT. Fixed cells were then washed with PBS, blocked in 5% BSA/PBS for 45 min followed by incubation with primary antibody in 5% BSA/PBS overnight at 4°C. Next day, the coverslips were washed with PBS and incubated with the secondary antibody for 1 h at RT. After washing with PBS, coverslips were mounted with Prolong Gold Antifade Mounting medium containing DAPI (ThermoFisher) and the coverslip edges were sealed with nail polish. After air-drying, images were acquired using Volocity software on an Axio Imager M2 microscope (Zeiss) equipped with an ORCA-ER camera (Hamamatsu) and a Plan-SPOCHROMAT 63×/1.4 oil objective. Images were processed using ImageJ software. For double immunostaining experiments (53BP1/Cyclin A) at least 100 nuclei were scored in each experiment. For the analysis of DAPI-positive chromosome bridges and lagging chromosomes, at least 50 anaphase cells were scored in each experiment. The primary antibodies used were: rabbit anti-53BP1 (ab36823; 1:500) and mouse anti-Cyclin A (sc-56299; 1:200). Secondary antibodies conjugated to Alexa Fluor 488 and Alexa Fluor 568 were used for immunodetection.

### Micronucleus formation

Cells grown on glass coverslips were treated with or without cordycepin. For quantification of micronuclei, cells were then treated with cytochalasin B (Sigma-Aldrich; 2 μg/ml for 16 h) to block cells in cytokinesis and fixed with PTEMF buffer for 10 min at RT. Slide mounting and imaging acquisition were performed as described above. At least 100 DAPI-stained binucleated cells were scored for the presence or absence of micronuclei in each experiment.

### Live-cell imaging

U2OS-GFP-m53BP1 cells (a gift from Dr Jiri Lukas) were seeded in Labtek chambers (Nunc, ThermoFisher) and after 24 h were transfected with either a SETX-targeting siRNA or a non-targeting negative control siRNA. After 36 h of siRNA transfection, the culture medium was changed to CO_2_-independent medium without phenol red (ThermoFisher) and live-cell imaging was carried out as described (37).

### Imagestream analysis

Cells were trypsinised and fixed in 70% ethanol for 1 hr at 4°C. They were then washed and resuspended at 2×10^7^ cells/ml in PBS containing 2% FBS. Cells were supplemented with 1 μg/ml propidium iodide (PI) and labelled for 15 minutes at room temperature. Flow cytometry and data collection was performed using the ImageStream®^X^ Mark II Imaging Flow Cytometer and Inspire acquisition software. Cells were imaged using channel 1 for bright field and fluorescent channel 5 (Bands 640-745 nm) and excitation laser 561 at 150-200mW. Image analysis was performed using the IDEAS software version 6.2. Single cells were filtered using a scatter plot of bright field aspect ratio and cell area. In-focus cells were filtered using a root mean square histogram with Gradient RMS > 50 within the bright field parameter. PI signal was inspected using the intensity channel 5 M05. Cells were inspected manually for micronuclei formation.

### Multiplex fluorescence *in situ* hybridization

Human 24-color multiplex fluorescence *in situ* hybridizations (M-FISH) were performed as described (105). At least 30 metaphases from each sample were karyotyped based on M-FISH classification and DAPI-banding pattern.

### Chromatin immunoprecipitation

Chromatin immunoprecipitation (ChIP) experiments were performed using the ChIP-IT Express Kit (Active Motif) as described (106). ChIP-derived DNA was purified using either the QIAquick PCR Purification Kit (Qiagen) or the Chromatin IP DNA Purification Kit (Active Motif). The following primary antibodies were used for ChIP experiments: rabbit anti-γH2AX (phospho S139; ab2893), rabbit anti-RPA (ab10359), rabbit anti-53BP1 (NB100-304), rabbit anti-CSB (A301-345A), rabbit anti-XPF (sc-28718), mouse anti-XPG (8H7) and mouse anti-RAD52 (sc-365341).

### DNA-RNA immunoprecipitation

DNA-RNA immunoprecipitation (DRIP) experiments were performed as described (8) with minor modifications. Briefly, nucleic acids were extracted using standard phenol-chloroform extraction and resuspended in nuclease-free water. Nucleic acids were digested with a cocktail of restriction enzymes (BsrGI, EcoRI, HindIII, SspI and XbaI) for 6 h at 37°C in the absence or presence of RNase H or RNase A (40 U, NEB). Digested nucleic acids were cleaned with phenol-chloroform extraction and resuspended in nuclease-free water. DNA-RNA hybrids (R-loops) were immunoprecipitated overnight at 4°C from 4 μg of total nucleic acids using 10 μg of S9.6 antibody in binding buffer (10 mM NaPO4 pH 7.0, 140 mM NaCl and 0.05% Triton X-100). Immunoprecipitated complexes were subsequently incubated with protein A/G sepharose (Abcam) for 2 h at 4°C, washed in binding buffer and eluted with elution buffer (50 mM Tris-HCl pH 8.0, 10 mM EDTA, 0.5% SDS). Proteinase K digestion was performed for 60 min at 55°C, followed by phenol-chloroform extraction and the DRIP-derived DNA pellet was resuspended in nuclease-free water.

### Quantitative real-time PCR

ChIP- and DRIP-derived DNA samples were subjected in triplicates (1.5 - 2 µl) to quantitative real-time PCR (qPCR) analysis on a CFX96 Real-Time Analyzer (Bio-Rad) using iQ SYBR Green Supermix reagent (Bio-Rad) and the gene-specific primers listed in the Table S7. The data were analyzed using the 2^−ΔΔCT^ method (107). Immunoprecipitated DNA was calculated as the percentage of DNA in the immunoprecipitates compared with input DNA. Fold enrichment of each target region was calculated as the ratio of the amounts of immunoprecipitated DNA estimated for the desired antibody versus control IgG.

### Gene expression array

Total RNA was purified using RNeasy Kit (Qiagen) from AOA2-mutant and respective control cells, according to the manufacturer’s instructions. Each condition was represented by three biological replicates. RNA samples were processed by services provided at the CRUK Manchester Institute using the NuGEN Amplification Kit (NuGEN) followed by hybridization to Affymetrix Human Gene 1.0 ST Array for gene expression. Arrays were scanned using the Affymetrix GeneArray 3000 7G scanner.

### Array comparative genomic hybridization

Genomic DNA from cultured human cells were isolated using the QIAamp DNA kit (Qiagen) and genomic DNA from MEFs and mouse tissue samples were extracted using the DNeasy Blood and Tissue Kit (Qiagen), according to the manufacturer’s recommendation. The purity and integrity of the DNA was verified by agarose gel electrophoresis with no RNA contamination observed. The Agilent SurePrint G3 Human CGH Microarrays 2×400K and the Agilent SurePrint G3 Mouse CGH Microarrays 1×1M were used for the experiments. Sample labeling, hybridization, washing and drying was carried out according to the manufacturer’s instructions (Agilent Technologies). Arrays were scanned using the NimbleGen MS 200 Microarray Scanner (Nimblegen-Roche).

### mRNA-sequencing

Total RNA was extracted from cultured human and mouse cells using an RNeasy Mini Kit (Qiagen) and included an on-column DNase treatment to eliminate contaminating genomic DNA. Samples were analysed on a 2100 Bioanalyzer (Agilent Technologies). All samples had an RNA Integrity Number (RIN) value greater than 8. The purified RNA was used for the preparation of poly A selected mRNA libraries using the TruSeq RNA sample preparation kit and sequenced on an Illumina HiSeq 4000 sequence analyser as paired-end 76 bp reads.

### RNAPII ChIP-sequencing

ChIP-sequencing (ChIP-seq) against RNAPII was performed as described (43). ChIP-derived DNA fragments were submitted for further manipulation by standard ChIP-seq library preparation techniques (Illumina) and Advanced Sequencing on an Illumina NextSeq 500 sequence analyzer as 75 bp single-end reads.

## Data availability

The microarray and sequencing data for this study was deposited at NCBI GEO under the accession number GEO: GSE143574. All other remaining data are available within the article and Supplementary Files, or available from the authors on request.

---

**Supplementary Figure S1.**
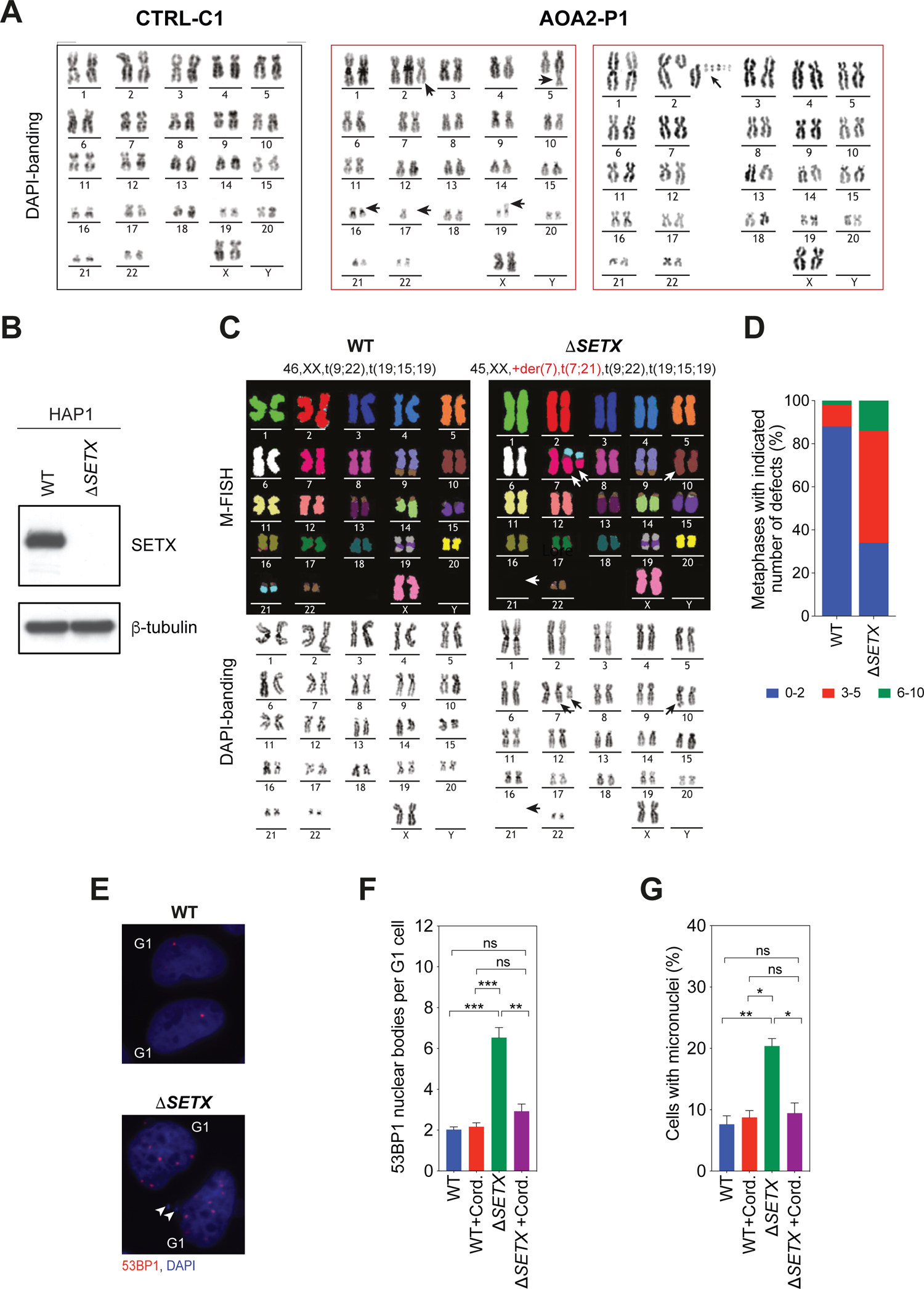
SETX deficiency promotes chromosome fragility. (A) DAP1-banding (gray-scale) images of M-FISH karyotypes shown in Fig 1F. Black arrows indicate chromosome aberrations. (B) Western blot analysis of HAP1 WT and Δ*SETX* cells made by CRISPR/Cas9-mediated gene targeting. β-tubulin was used as a loading control. (C) M-FISH and DAPI-banding of metaphase spreads from HAP1 WT and Δ*SETX* cells showing deletions, translocations and chromosome fragility. Representative karyotypes are shown. Arrows indicate chromosome aberrations. (D) Quantification of metaphases with indicated number of aberrations, as in (C). 30 metaphases were analysed per condition. (E) Immunostaining of HAP1 WT and Δ*SETX* cells with 53BP1 (red) antibody. Nuclear DNA was stained with DAPI (blue). Representative images of G1-phase cells are shown. White arrowheads indicate micronuclei. (F) Quantification of 53BP1 nuclear bodies in G1 cells, as in (E). Cells were treated with or without cordycepin (Cord.). (G) Quantification of cells with indicated number of micronuclei, as in (E). Cells were treated with or without cordycepin. Data are representative of three independent experiments. *P<0.05, **P<0.01 and ***P<0.001 by Mann-Whitney test. P≥0.05 is considered not significant (ns). Error bars represent s.e.m.

**Supplementary Figure S2.**
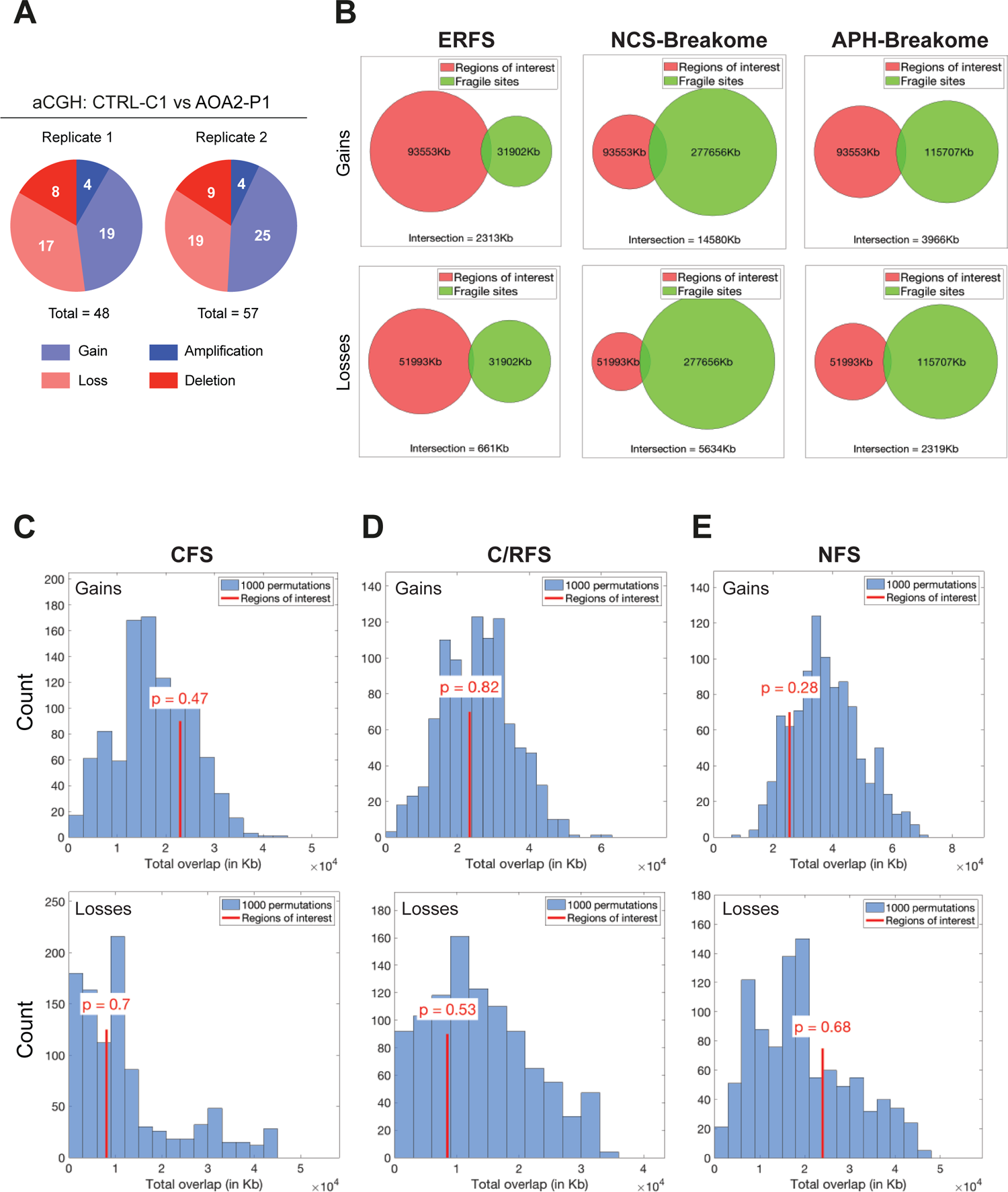
Overlap analysis of CNCs identified in AOA2-P1 fibroblasts by array comparative genomic hybridization. (A) Pie chart representations of CNCs identified in the aCGH experiments described in Fig 2A and B. Data from two independent experiments (replicates 1 and 2) are shown. (B) Venn diagram of AOA2-P1 fibroblast gain or loss regions (red) and fragile sites (green) as in Fig 2C-E. Total genomic area and intersected area are reported. (C) Top: Histogram showing overlaps between 1000 permuted AOA2-P1 fibroblast gain regions with CFSs. Bottom: Histogram showing the overlaps between 1000 permuted AOA2-P1 fibroblast loss regions with CFSs. Red line indicates the degree of overlap (in kb). P values for the overlap compared to permutations are indicated, with P<0.05 considered a significant enrichment/depletion. (D) As (C) showing the overlaps between AOA2-P1 fibroblast gain and loss regions with the C/RFS. (E) As (C) showing the overlaps between AOA2-P1 fibroblast gain and loss regions with the NFS.

**Supplementary Figure S3.**
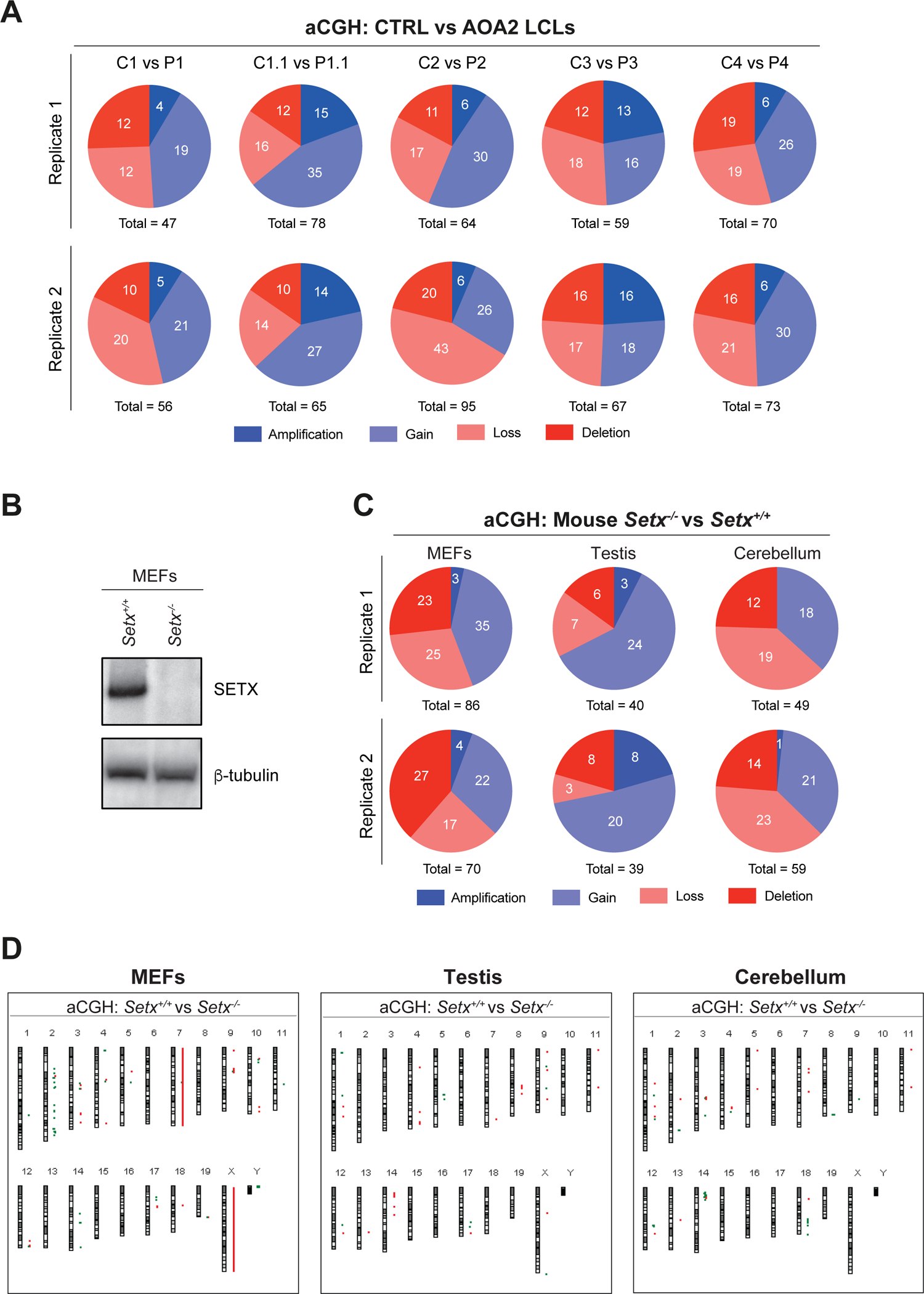
Genomic aberrations associated with AOA2 LCLs and mouse *Setx* k/o. (A) Pie chart representations of CNCs detected in two independent aCGH experiments (replicates 1 and 2) with genomic DNA from control (C1-C4 and C1.1) and AOA2 (P1-P4 and P1.1) LCLs. (B) Western blot analysis of *Setx^+/+^* and *Setx^-/-^* MEFs. β-tubulin was used as a loading control. (C) As (A) but with genomic DNA from *Setx^+/+^* and *Setx^-/-^* MEFs, testis and cerebellum. (D) Ideograms indicating the chromosomal locations of CNCs. Regions of gain and loss are indicated with red and green bars, respectively.

**Supplementary Figure S4.**
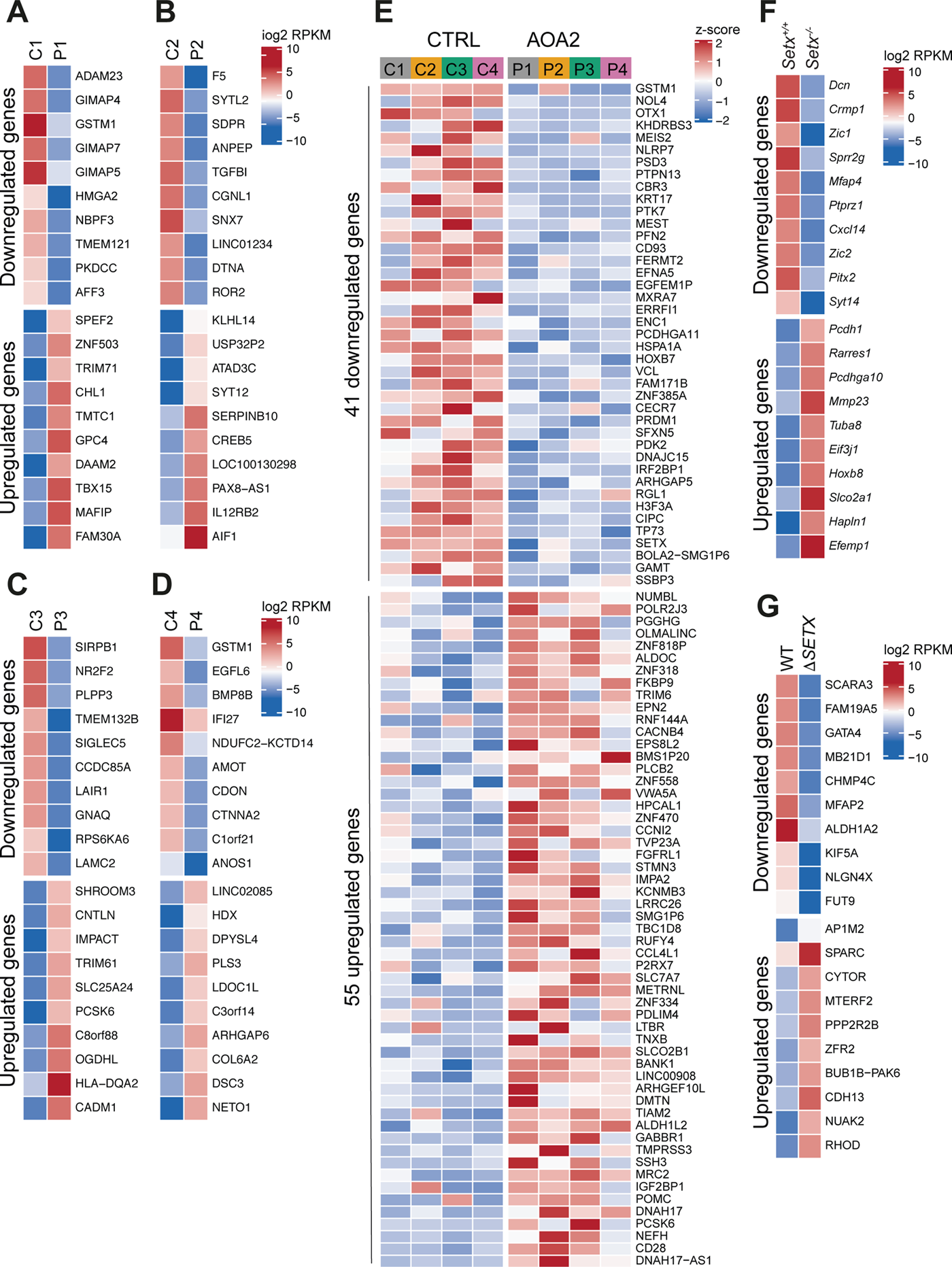
Differentially expressed genes identified in SETX-deficient human and mouse cells. (A) Heatmap of gene expression abundance across P1 and C1 LCLs. The top 10 most downregulated and 10 most upregulated genes, based on a fold change of normalised read count, are shown. Data are presented as log2(RPKM+0.01). (B) As (A), using RNA-seq analyses from C2 and P2 LCLs. (C) As (A), using RNA-seq analyses from C3 and P3 LCLs. (D) As (A), using RNA-seq analyses from C4 and P4 LCLs. (E) Heatmap of gene expression abundance, highlighting differences in expression between AOA2 and paired control LCLs from DESeq2 analysis (FDR≤0.05). All the downregulated (41) and upregulated (55) genes based on fold change are presented. Data are rlog transformed to stabilise variance and normalise with respect to library size, then scaled per-gene using a z-score. (F) As (A), using RNA-seq analyses from *Setx*^+/+^ and *Setx*^-/-^ mouse MEFs. (G) As (A), using RNA-seq analyses from HAP1 WT and Δ*SETX* cells.

**Supplementary Figure S5.**
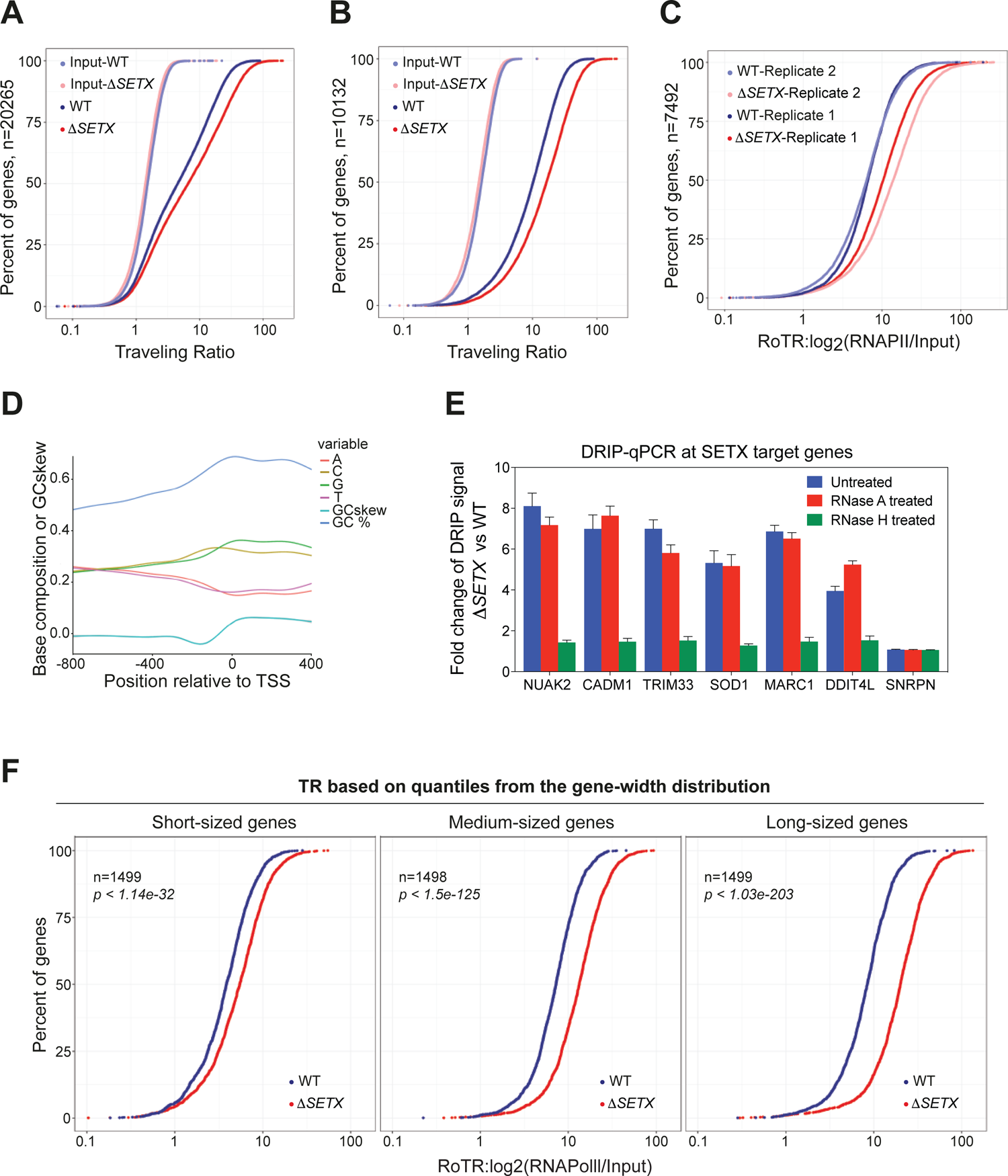
Transcription stress analysis in HAP1 WT and Δ*SETX* cells. (A) RNAPII traveling ratio (TR) distribution of all genes (n=20265) in HAP1 WT and Δ*SETX* cells, along with input controls. The y-axis indicates percent of all genes. Higher TR values indicate a higher degree of RNAPII pausing. (B) As (A), except TR was analysed only for genes with RNAPII peak over TSS (n=10132). (C) As Fig 4C, except that cumulative curves of RNAPII RoTR from two independent ChIP-seq experiments are shown. (D) Nucleotide frequencies in the [-800, +400] region around the TSS of 7492 genes identified in the RoTR analysis. GC skew and GC content (GC%) in the region are shown. (E) DRIP-qPCR assays were carried out using HAP1 WT and Δ*SETX* cells, either untreated or treated with RNase A or RNase H. The SETX-target gene promoter and negative control (SNRPN) regions were analysed. Data represent the mean ± s.e.m of three independent experiments. (F) As Fig 4E, except that Ensembl genes are stratified into short/medium/long groupings based on the quantiles from the gene-width distributions. ‘short’ = shortest 20%, ‘medium’ = middle 40-60% of widths and ‘long’ = longest 20%.

**Supplementary Figure S6.**
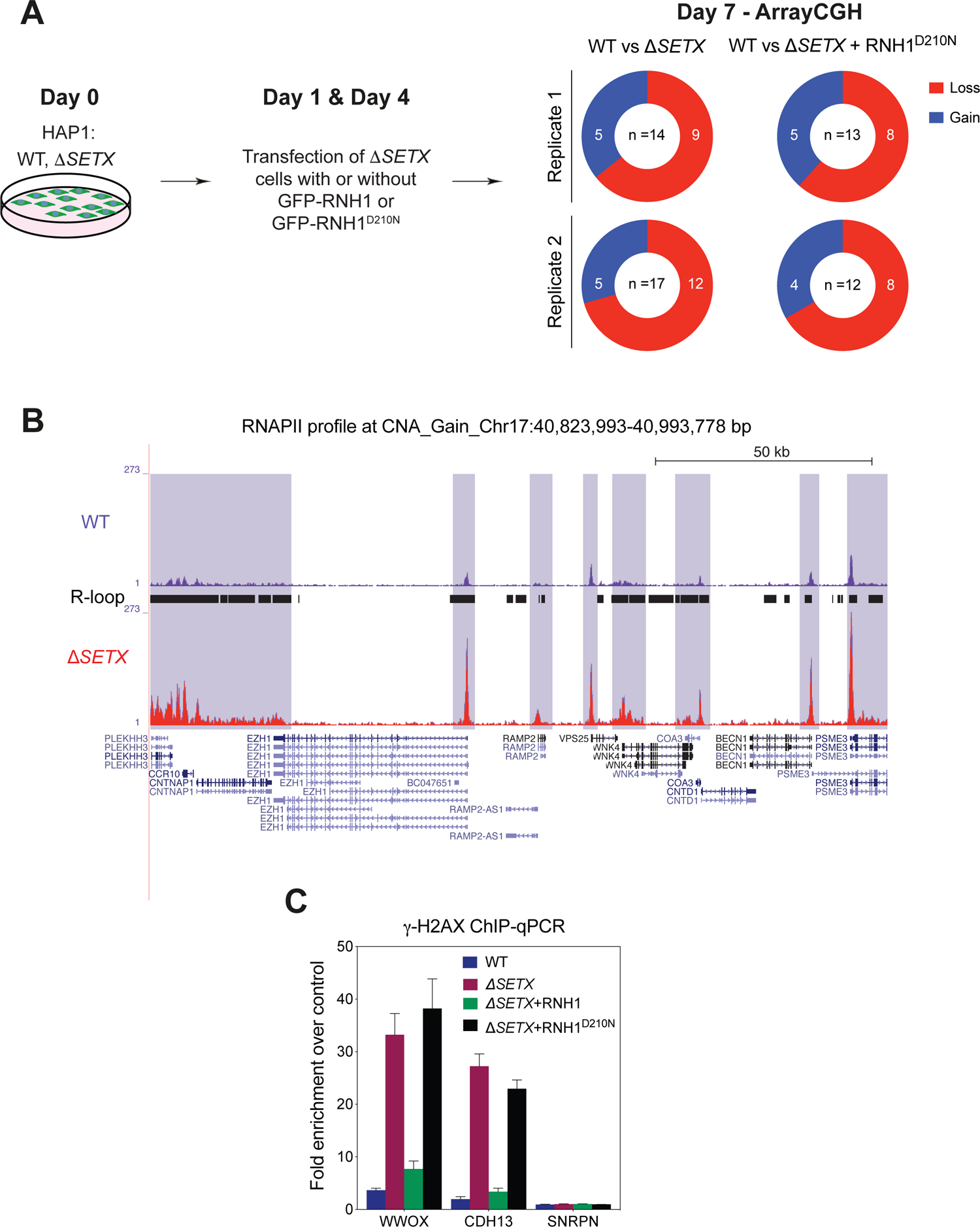
Transcription stress causes chromosome instability in the absence of SETX. (A) Schematic of aCGH experiments carried out with HAP1 WT and Δ*SETX* cells. On day 1 and day 4 post seeding, Δ*SETX* cells were transfected with or without GFP-RNaseH1 (RNH1) or GFP-RNH1^D210N^. On day 7, aCGH was performed with WT vs WT, WT vs Δ*SETX*, WT vs Δ*SETX* + RNH1 and WT vs Δ*SETX* + RNH1^D210N^. Donut chart representations of CNCs identified in the aCGH experiments described in Fig 5A and D. Data from two independent experiments (replicates 1 and 2) are shown. (B) Representative genome browser screenshot of RNAPII pausing over the indicated genomic region that is amplified in Δ*SETX* (red) compared to WT (blue) cells. R-loops are shown in black. All genes in the region are shown below. Scale bar, 50kb. (C) ChIP-qPCR analyses at *WWOX*, *CDH13 and SNRPN* (negative control) genes using a γ-H2AX antibody was carried out with cross-linked chromatin from HAP1 WT and Δ*SETX* cells. Fold enrichment was calculated as a ratio of γ-H2AX antibody signal versus control IgG. Data represent the mean ± s.e.m of three independent experiments.

**Supplementary Figure S7.**
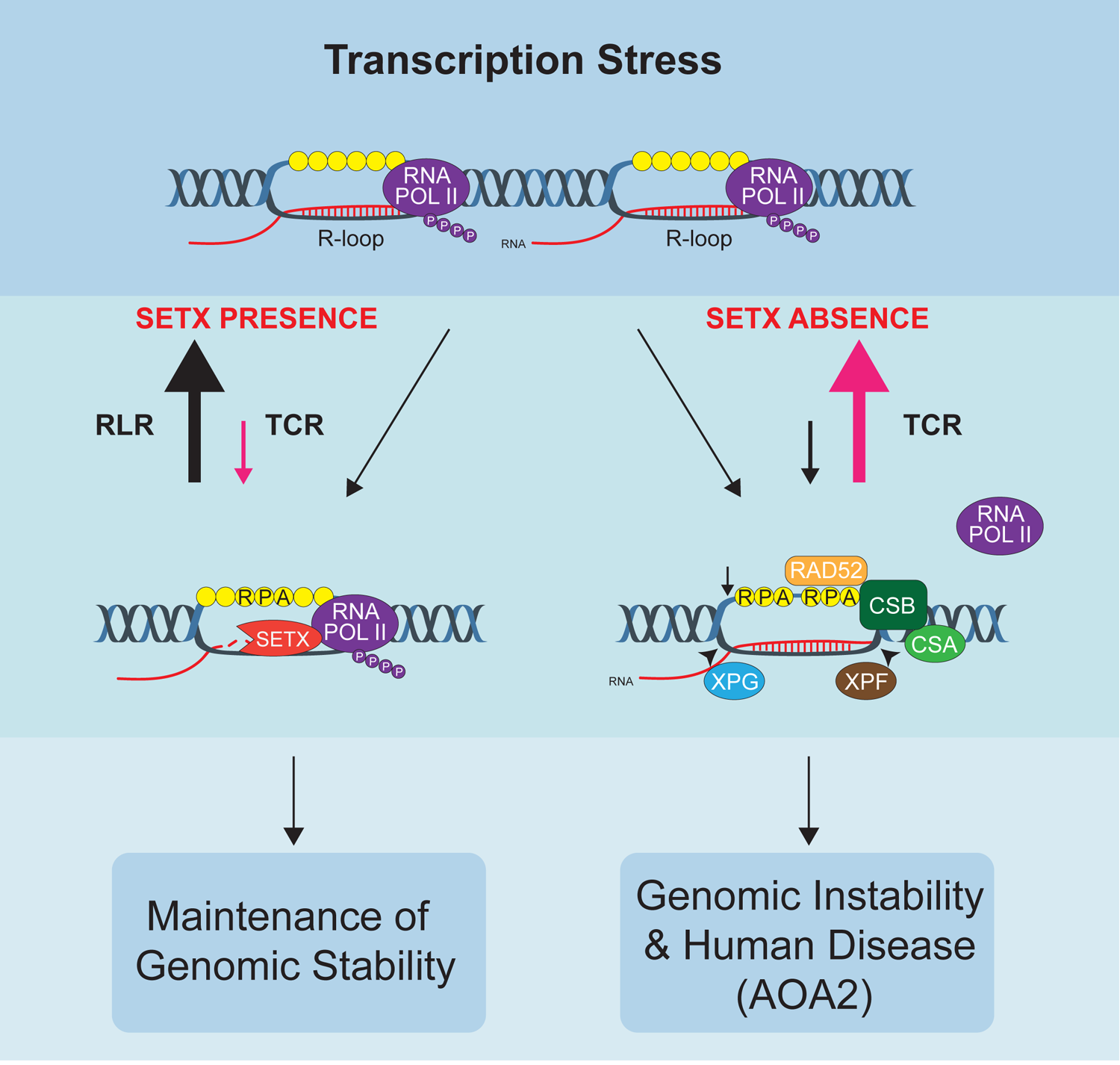
Model for the role of SETX in response to transcription stress (RNAPII pausing, ROS damage). Senataxin facilitates R-loop repair (RLR) by promoting error-free removal of R-loops at transcription sites, particularly near regions of RNAPII pausing, and thereby protects against genome stability. In the absence of SETX, CSB recognises stalled RNAPII and transcription-coupled repair (TCR) proteins/recombination factors (e.g. RAD52) are recruited to resolve and repair transcription bubbles containing R-loops, but at the expense of faithful maintenance of the genome.

## REFERENCES

1. Gaillard, H. and Aguilera, A. (2016) Transcription as a threat to genome integrity. Annu. Rev. Biochem., 85, 291–317.

2. Gomez-Gonzalez, B. and Aguilera, A. (2019) Transcription-mediated replication hindrance: a major driver of genome instability. Genes Dev., 33, 1008–1026.

3. Lang, K.S. and Merrikh, H. (2018) The clash of macromolecular titans: replication-transcription conflicts in bacteria. Annu. Rev. Microbiol., 72, 71–88.

4. Skourti-Stathaki, K., Kamieniarz-Gdula, K. and Proudfoot, N.J. (2014) R-loops induce repressive chromatin marks over mammalian gene terminators. Nature, 516, 436–439.

5. Crossley, M.P., Bocek, M. and Cimprich, K.A. (2019) R-Loops as cellular regulators and genomic threats. Mol. Cell, 73, 398–411.

6. Chen, L., Chen, J.Y., Zhang, X., Gu, Y., Xiao, R., Shao, C., Tang, P., Qian, H., Luo, D., Li, H. et al. (2017) R-ChIP using inactive RNase H reveals dynamic coupling of R-loops with transcriptional pausing at gene promoters. Mol. Cell, 68, 745–757.

7. Ginno, P.A., Lim, Y.W., Lott, P.L., Korf, I. and Chedin, F. (2013) GC skew at the 5’ and 3’ ends of human genes links R-loop formation to epigenetic regulation and transcription termination. Genome Res., 23, 1590–1600.

8. Ginno, P.A., Lott, P.L., Christensen, H.C., Korf, I. and Chedin, F. (2012) R-loop formation is a distinctive characteristic of unmethylated human CpG island promoters. Mol. Cell, 45, 814–825.

9. Skourti-Stathaki, K., Proudfoot, N.J. and Gromak, N. (2011) Human Senataxin resolves RNA/DNA hybrids formed at transcriptional pause sites to promote Xrn2-dependent transcription. Mol. Cell, 42, 794–805.

10. El Hage, A., French, S.L., Beyer, A.L. and Tollervey, D. (2010) Loss of Topoisomerase I leads to R-loop-mediated transcriptional blocks during ribosomal RNA synthesis. Genes Dev., 24, 1546–1558.

11. Chan, Y.A., Aristizabal, M.J., Lu, P.Y., Luo, Z., Hamza, A., Kobor, M.S., Stirling, P.C. and Hieter, P. (2014) Genome-wide profiling of yeast DNA:RNA hybrid prone sites with DRIP-chip. PLoS Genet., 10, e1004288.

12. Ohle, C., Tesorero, R., Schermann, G., Dobrev, N., Sinning, I. and Fischer, T. (2016) Transient RNA-DNA hybrids are required for efficient double-strand break repair. Cell, 167, 1001–1013.

13. Baker, T.A. and Kornberg, A. (1988) Transcriptional activation of initiation of replication from the *E. coli* chromosomal origin: an RNA-DNA hybrid near oriC. Cell, 55, 113–123.

14. Xu, B. and Clayton, D.A. (1996) RNA-DNA hybrid formation at the human mitochondrial heavy-strand origin ceases at replication start sites: an implication for RNA-DNA hybrids serving as primers. EMBO J., 15, 3135–3143.

15. Yu, K., Chedin, F., Hsieh, C.L., Wilson, T.E. and Lieber, M.R. (2003) R-loops at immunoglobulin class switch regions in the chromosomes of stimulated B cells. Nat. Immunol., 4, 442–451.

16. Sanz, L.A., Hartono, S.R., Lim, Y.W., Steyaert, S., Rajpurkar, A., Ginno, P.A., Xu, X. and Chedin, F. (2016) Prevalent, dynamic and conserved R-loop structures associate with specific epigenomic signatures in mammals. Mol. Cell, 63, 167–178.

17. Garcia-Muse, T. and Aguilera, A. (2019) R-loops: from physiological to pathological roles. Cell, 179, 604–618.

18. Cerritelli, S.M. and Crouch, R.J. (2009) Ribonuclease H: the enzymes in eukaryotes. FEBS J., 276, 1494–1505.

19. Mischo, H., Gómez-González, B., Grezechnik, P., Rondón, A.G., Wei, W., Steinmetz, L.M., Aguilera, A. and Proudfoot, N.J. (2011) Yeast Sen1 helicase protects the genome from transcription-associated instability. Mol. Cell, 41, 21–32.

20. Wahba, L., Amon, J.D., Koshland, D. and Vuica-Ross, M. (2011) RNase H and multiple RNA biogenesis factors cooperate to prevent RNA:DNA hybrids from generating genome instability. Mol. Cell, 44, 978–988.

21. Sollier, J., Stork, C.T., Garcia-Rubio, M.L., Paulsen, R.D., Aguilera, A. and Cimprich, K.A. (2014) Transcription-coupled nucleotide excision repair factors promote R-loop-induced genome instability. Mol. Cell, 56, 777–785.

22. Perez-Calero, C., Bayona-Feliu, A., Xue, X., Barroso, S.I., Munoz, S., Gonzalez-Basallote, V.M., Sung, P. and Aguilera, A. (2020) UAP56/DDX39B is a major cotranscriptional RNA-DNA helicase that unwinds harmful R loops genome-wide. Genes Dev., 34, 898–912.

23. Moreira, M.C., Klur, S., Watanabe, M., Nemeth, A.H., Le Ber, I., Moniz, J.C., Tranchant, C., Aubourg, P., Tazir, M., Schols, L. et al. (2004) Senataxin, the ortholog of a yeast RNA helicase, is mutant in ataxia-ocular apraxia 2. Nat. Genet., 36, 225–227.

24. Chen, Y.Z., Bennett, C.L., Huynh, H.M., Blair, I.P., Puls, I., Irobi, J., Dierick, I., Abel, A., Kennerson, M.L., Rabin, B.A. et al. (2004) DNA/RNA helicase gene mutations in a form of juvenile amyotrophic lateral sclerosis (ALS4). Am. J. Hum. Genet., 74, 1128–1135.

25. Lavin, M.F., Yeo, A.J. and Becherel, O.J. (2013) Senataxin protects the genome: Implications for neurodegeneration and other abnormalities. Rare Dis., 1, e25230.

26. Le Ber, I., Bouslam, N., Rivaud-Pechoux, S., Guimaraes, J., Benomar, A., Chamayou, C., Goizet, C., Moreira, M.C., Klur, S., Yahyaoui, M. et al. (2004) Frequency and phenotypic spectrum of ataxia with oculomotor apraxia 2: a clinical and genetic study in 18 patients. Brain, 127, 759–767.

27. De Amicis, A., Piane, M., Ferrari, F., Fanciulli, M., Delia, D. and Chessa, L. (2011) Role of senataxin in DNA damage and telomeric stability. DNA Rep., 10, 199–209.

28. Roda, R.H., Rinaldi, C., Singh, R.K., Schindler, A.B. and Blackstone, C. (2014) Ataxia with oculomotor apraxia type 2 fibroblasts exhibit increased susceptibility to oxidative DNA damage. J. Clin. Neurosci., 21, 1627–1631.

29. Suraweera, A., Becherel, O.J., Chen, P., Rundle, N., Woods, R.A., Nakamura, J., Gatei, M., Criscuolo, C., Filla, A., Chessa, L. et al. (2007) Senataxin, defective in ataxia oculomotor apraxia 2, is involved in the defence against oxidative DNA damage. J. Cell. Biol., 177, 969–979.

30. Becherel, O.J., Sun, J., Yeo, A.J., Nayler, S., Fogel, B.L., Gao, F., Coppola, G., Criscuolo, C., De Michele, G., Wolvetang, E. et al. (2015) A new model to study neurodegeneration in ataxia oculomotor apraxia type 2. Hum. Mol. Genet., 24, 5759–5774.

31. Becherel, O.J., Yeo, A.J., Stellati, A., Heng, E.Y.H., Luff, J.E., Suraweera, A.M., Woods, R.A., Fleming, J., Carrie, D., McKinney, K. et al. (2013) Senataxin plays an essential role with DNA damage response proteins in meiotic recombination and gene silencing. PLoS Genet., 9, e1003435.

32. Grunseich, C., Wang, I.X., Watts, J.A., Burdick, J.T., Guber, R.D., Zhu, Z., Bruzel, A., Lanman, T., Chen, K., Schindler, A.B. et al. (2018) Senataxin mutation reveals how R-loops promote transcription by blocking DNA methylation at gene promoters. Mol. Cell, 69, 426–437.

33. Hatchi, E., Skourti-Stathaki, K., Ventz, S., Pinello, L., Yen, A., Kamieniarz-Gdula, K., Dimitrov, S.D., Pathania, S., McKinney, K.M., Eaton, M.L. et al. (2015) BRCA1 recruitment to transcriptional pause sites is required for R-loop-driven DNA damage repair. Mol. Cell, 57, 636–647.

34. Suraweera, A., lim, Y., Woods, R.A., Birrell, G.W., Nasim, T., Becherel, O.J. and Lavin, M.F. (2009) Functional role for Senataxin, defective in ataxia oculomotor apraxia type 2, in transcription regulation. Human Mol. Genet., 18, 3384–3396.

35. Zhao, D.Y., Gish, G., Braunschweig, U., Li, Y., Ni, Z., Schmitges, F.W., Zhong, G., Liu, K., Li, W., Moffat, J. et al. (2016) SMN and symmetric arginine dimethylation of RNA polymerase II C-terminal domain control termination. Nature, 529, 48–53.

36. Richard, P. and Manley, J.L. (2014) SETX sumoylation: A link between DNA damage and RNA surveillance disrupted in AOA2. Rare Dis., 2, e27744.

37. Yüce, O. and West, S.C. (2013) Senataxin, defective in the neurodegenerative disorder Ataxia with Oculomotor Apraxia 2, lies at the interface of transcription and the DNA damage response. Mol. Cell. Biol., 33, 406–417.

38. Andrews, A.M., McCartney, H.J., Errington, T.M., D’Andrea, A.D. and Macara, I.G. (2018) A senataxin-associated exonuclease SAN1 is required for resistance to DNA interstrand cross-links. Nat Commun, 9, 2592.

39. Cohen, S., Puget, N., Lin, Y.L., Clouaire, T., Aguirrebengoa, M., Rocher, V., Pasero, P., Canitrot, Y. and Legube, G. (2018) Senataxin resolves RNA:DNA hybrids forming at DNA double-strand breaks to prevent translocations. Nat. Commun., 9, 533.

40. Kannan, A., Bhatia, K., Branzei, D. and Gangwani, L. (2018) Combined deficiency of Senataxin and DNA-PKcs causes DNA damage accumulation and neurodegeneration in spinal muscular atrophy. Nucl. Acids Res., 46, 8326–8346.

41. Yeo, A.J., Becherel, O.J., Luff, J.E., Graham, M.E., Richard, D. and Lavin, M.F. (2015) Senataxin controls meiotic silencing through ATR activation and chromatin remodeling. Cell Discov., 1, 15025.

42. Miller, M.S., Rialdi, A., Ho, J.S., Tilove, M., Martinez-Gil, L., Moshkina, N.P., Peralta, Z., Noel, J., Melegari, C., Maestre, A.M. et al. (2015) Senataxin suppresses the antiviral transcriptional response and controls viral biogenesis. Nat. Immunol., 16, 485–494.

43. Saponaro, M., Kantidakis, T., Mitter, R., Kelly, G.P., Heron, M., Williams, H.R., Söding, J., Stewart, A.F. and Svejstrup, J.Q. (2014) RECQL5 controls transcript elongation and suppresses genome instability associated with transcription stress. Cell, 157, 1037–1049.

44. Kantidakis, T., Saponaro, M., Mitter, R., Horswell, S., Kranz, A., Boeing, S., Aygun, O., Kelly, G.P., Matthews, N., Stewart, A. et al. (2016) Mutation of cancer driver MLL2 results in transcription stress and genome instability. Genes Dev., 30, 408–420.

45. Barlow, J.H., Faryabi, R.B., Callen, E., Wong, N., Malhowski, A., Chen, H.T., Gutierrez-Cruz, G., Sun, H.W., McKinnon, P., Wright, G. et al. (2013) Identification of early replicating fragile sites that contribute to genome instability. Cell, 152, 620–632.

46. Crosetto, N., Mitra, A., Silva, M.J., Bienko, M., Dojer, N., Wang, Q., Karaca, E., Chiarle, R., Skrzypczak, M., Ginalski, K. et al. (2013) Nucleotide-resolution DNA double-strand break mapping by next-generation sequencing. Nat. Methods, 10, 361–365.

47. Bignell, G.R., Greenman, C.D., Davies, H., Butler, A.P., Edkins, S., Andrews, J.M., Buck, G., Chen, L., Beare, D., Latimer, C. et al. (2010) Signatures of mutation and selection in the cancer genome. Nature, 463, 893–898.

48. Fungtammasan, A., Walsh, E., Chiaromonte, F., Eckert, K.A. and Makova, K.D. (2012) A genome-wide analysis of common fragile sites: what features determine chromosomal instability in the human genome? Genome Res., 22, 993–1005.

49. Helmrich, A., Stout-Weider, K., Hermann, K., Schrock, E. and Heiden, T. (2006) Common fragile sites are conserved features of human and mouse chromosomes and relate to large active genes. Genome Res., 16, 1222–1230.

50. Wei, P.C., Chang, A.N., Kao, J., Du, Z., Meyers, R.M., Alt, F.W. and Schwer, B. (2016) Long neural genes harbor recurrent DNA break clusters in neural stem/progenitor cells. Cell, 164, 644–655.

51. Wei, P.C., Lee, C.S., Du, Z., Schwer, B., Zhang, Y., Kao, J., Zurita, J. and Alt, F.W. (2018) Three classes of recurrent DNA break clusters in brain progenitors identified by 3D proximity-based break joining assay. Proc. Natl. Acad. Sci .U. S. A., 115, 1919–1924.

52. Fogel, B.L., Cho, E., Wahnich, A., Gao, F., Becherel, O.J., Wang, X., Fike, F., Chen, L., Criscuolo, C., De Michele, G. et al. (2014) Mutation of Senataxin alters disease-specific transcriptional networks in patients with ataxia with oculomotor apraxia type 2. Hum. Mol. Genet., 23, 4758–4769.

53. Rahl, P.B., Lin, C.Y., Seila, A.C., Flynn, R.A., McCuine, S., Burge, C.B., Sharp, P.A. and Young, R.A. (2010) c-Myc regulates transcriptional pause release. Cell, 141, 432–445.

54. Helmrich, A., Ballarino, M. and Tora, L. (2011) Collisions between replication and transcription complexes cause common fragile site instability at the longest human genes. Mol. Cell, 44, 966–977.

55. Tuduri, S., Crabbé, L., Conti, C., Tourierre, H., Holtgreve-Grez, H., Jauch, A., Pantesco, V., De Vos, J., Thomas, A., Theillet, C. et al. (2009) Topoisomerase I suppresses genomic instability by preventing interference between replication and transcription. Nat. Cell Biol., 11, 1315–1324.

56. Sollier, J. and Cimprich, K.A. (2015) Breaking bad: R-loops and genome integrity. Trends Cell Biol., 25, 514–522.

57. Aguilera, A. and Gomez-Gonzalez, B. (2017) DNA-RNA hybrids: the risks of DNA breakage during transcription. Nat. Struct. Mol. Biol., 24, 439–443.

58. Nilson, K.A., Lawson, C.K., Mullen, N.J., Ball, C.B., Spector, B.M., Meier, J.L. and Price, D.H. (2017) Oxidative stress rapidly stabilizes promoter-proximal paused Pol II across the human genome. Nucl. Acids Res., 45, 11088–11105.

59. Yasuhara, T., Kato, R., Hagiwara, Y., Shiotani, B., Yamauchi, M., Nakada, S., Shibata, A. and Miyagawa, K. (2018) Human RAD52 promotes XPG-mediated R-loop processing to initiate transcription-associated homologous recombination repair. Cell, 175, 558–570.

60. Teng, Y., Yadav, T., Duan, M., Tan, J., Xiang, Y., Gao, B., Xu, J., Liang, Z., Liu, Y., Nakajima, S. et al. (2018) ROS-induced R loops trigger a transcription-coupled but BRCA1/2-independent homologous recombination pathway through CSB. Nat. Commun., 9, 4115.

61. Lemay, J.F., Larochelle, M., Marguerat, S., Atkinson, S., Bahler, J. and Bachand, F. (2014) The RNA exosome promotes transcription termination of backtracked RNA polymerase II. Nat. Struct. Mol. Biol., 21, 919–926.

62. Sheridan, R.M., Fong, N., D’Alessandro, A. and Bentley, D.L. (2019) Widespread backtracking by RNA Pol II is a major effector of gene activation, 5′ pause release, termination, and transcription elongation rate. Mol Cell, 73, 107-118.

63. Zatreanu, D., Han, Z., Mitter, R., Tumini, E., Williams, H., Gregersen, L., Dirac-Svejstrup, A.B., Roma, S., Stewart, A., Aguilera, A. et al. (2019) Elongation factor TFIIS prevents transcription stress and R-loop accumulation to maintain genome stability. Mol. Cell, 76, 57–69.

64. Porrua, O. and Libri, D. (2013) A bacterial-like mechanism for transcription termination by the Sen1p helicase in budding yeast. Nat. Struct. Mol. Biol., 20, 884–891.

65. Steinmetz, E.J., Warren, C.D., Kuehner, J.N., Panbehi, B., Ansari, A.Z. and Brow, D.A. (2006) Genome-wide distribution of yeast RNA polymerase II and its control by Sen1 helicase. Mol. Cell, 24, 735–746.

66. Ursic, D., Himmel, K.L., Gurley, K.A., Webb, F. and Culbertson, M.R. (1997) The yeast *SEN1* gene is required for the processing of diverse RNA classes. Nucl. Acids Res., 25, 4778–4785.

67. Connelly, S. and Manley, J.L. (1988) A functional mRNA polyadenylation signal is required for transcription termination by RNA polymerase II. Genes Dev., 2, 440–452.

68. Kim, M., Krogan, N.J., Vasiljeva, L., Rando, O.J., Nedea, E., Greenblatt, J.F. and Buratowski, S. (2004) The yeast Rat1 exonuclease promotes transcription termination by RNA polymerase II. Nature, 432, 517–522.

69. Proudfoot, N.J. (2011) Ending the message: poly(A) signals then and now. Genes Dev., 25, 1770–1782.

70. Proudfoot, N.J. (2016) Transcriptional termination in mammals: Stopping the RNA polymerase II juggernaut. Science, 352, aad9926.

71. West, S., Gromak, N. and Proudfoot, N.J. (2004) Human 5’ --> 3’ exonuclease Xrn2 promotes transcription termination at co-transcriptional cleavage sites. Nature, 432, 522–525.

72. Gregersen, L.H., Mitter, R., Ugalde, A.P., Nojima, T., Proudfoot, N.J., Agami, R., Stewart, A. and Svejstrup, J.Q. (2019) SCAF4 and SCAF8, mRNA anti-terminator proteins. Cell, 177, 1797–1813.

73. Gomez-Gonzalez, B., Garcia-Rubio, M., Bermejo, R., Gaillard, H., Shirahige, K., Marin, A., Foiani, M. and Aguilera, A. (2011) Genome-wide function of THO/TREX in active genes prevents R-loop-dependent replication obstacles. EMBO J., 30, 3106–3119.

74. Stirling, P.C., Chan, Y.A., Minaker, S.W., Aristizabal, M.J., Barrett, I., Sipahimalani, P., Kobor, M.S. and Hieter, P. (2012) R-loop-mediated genome instability in mRNA cleavage and polyadenylation mutants. Genes Dev., 26, 163–175.

75. Grzechnik, P., Tan-Wong, S.M. and Proudfoot, N.J. (2014) Terminate and make a loop: regulation of transcriptional directionality. Trends Biochem. Sci., 39, 319–327.

76. Madabhushi, R., Pan, L. and Tsai, L.H. (2014) DNA damage and its links to neurodegeneration. Neuron, 83, 266–282.

77. McKinnon, P.J. (2017) Genome integrity and disease prevention in the nervous system. Genes Dev., 31, 1180–1194.

78. Bassuk, A.G., Chen, Y.Z., Batish, S.D., Nagan, N., Opal, P., Chance, P.F. and Bennett, C.L. (2007) In cis autosomal dominant mutation of Senataxin associated with tremor/ataxia syndrome. Neurogenetics, 8, 45–49.

79. Høyer, H., Braathen, G.J., Busk, O.L., Holla, O.L., Svendsen, M., Hilmarsen, H.T., Strand, L., Skjelbred, C.F. and Russell, M.B. (2014) Genetic diagnosis of Charcot-Marie-Tooth disease in a population by next-generation sequencing. Biomed. Res. Int., 2014, 210401.

80. Rudnik-Schoneborn, S., Arning, L., Epplen, J.T. and Zerres, K. (2012) *SETX* gene mutation in a family diagnosed autosomal dominant proximal spinal muscular atrophy. Neuromuscul. Disord., 22, 258–262.

81. Criscuolo, C., Chessa, L., Di Giandomenico, S., Mancini, P., Sacca, F., Grieco, G.S., Piane, M., Barbieri, F., De Michele, G., Banfi, S. et al. (2006) Ataxia with oculomotor apraxia type 2: a clinical, pathologic, and genetic study. Neurology, 66, 1207–1210.

82. Anheim, M., Monga, B., Fleury, M., Charles, P., Barbot, C., Salih, M., Delaunoy, J.P., Fritsch, M., Arning, L., Synofzik, M. et al. (2009) Ataxia with oculomotor apraxia type 2: clinical, biological and genotype/phenotype correlation study of a cohort of 90 patients. Brain, 132, 2688–2698.

83. Nanetti, L., Cavalieri, S., Pensato, V., Erbetta, A., Pareyson, D., Panzeri, M., Zorzi, G., Antozzi, C., Moroni, I., Gellera, C. et al. (2013) SETX mutations are a frequent genetic cause of juvenile and adult onset cerebellar ataxia with neuropathy and elevated serum alpha-fetoprotein. Orphanet. J. Rare Dis., 8, 123.

84. Airoldi, G., Guidarelli, A., Cantoni, O., Panzeri, C., Vantaggiato, C., Bonato, S., Grazia D’Angelo, M., Falcone, S., De Palma, C., Tonelli, A. et al. (2010) Characterization of two novel *SETX* mutations in AOA2 patients reveals aspects of the pathophysiological role of senataxin. Neurogenetics, 11, 91–100.

85. Wang, Y., Chakravarty, P., Ranes, M., Kelly, G.P., Brooks, P.J., Neilan, E., Stewart, A., Schiavo, G. and Svejstrup, J.Q. (2014) Dysregulation of gene expression as a cause of Cockayne syndrome neurological disease. Proc. Natl. Acad. Sci .U. S. A., 111, 14454–14459.

86. Carette, J.E., Raaben, M., Wong, A.C., Herbert, A.S., Obernosterer, G., Mulherkar, N., Kuehne, A.I., Kranzusch, P.J., Griffin, A.M., Ruthel, G. et al. (2011) Ebola virus entry requires the cholesterol transporter Niemann-Pick C1. Nature, 477, 340–343.

87. Bekker-Jensen, S., Lukas, C., Melander, F., Bartek, J. and Lukas, J. (2005) Dynamic assembly and sustained retention of 53BP1 at the sites of DNA damage are controlled by Mdc1/NFBD1. J. Cell Biol., 170, 201–211.

88. Suzuki, Y., Holmes, J.B., Cerritelli, S.M., Sakhuja, K., Minczuk, M., Holt, I.J. and Crouch, R.J. (2010) An upstream open reading frame and the context of the two AUG codons affect the abundance of mitochondrial and nuclear RNase H1. Mol. Cell. Biol., 30, 5123–5134.

89. Hu, Z., Zhang, A., Storz, G., Gottesman, S. and Leppla, S.H. (2006) An antibody-based microarray assay for small RNA detection. Nucleic Acids Res, 34, e52.

90. Quinlan, A.R. and Hall, I.M. (2010) BEDTools: a flexible suite of utilities for comparing genomic features. Bioinformatics, 26, 841–842.

91. Li, H., Handsaker, B., Wysoker, A., Fennell, T., Ruan, J., Homer, N., Marth, G., Abecasis, G., Durbin, R. and Genome Project Data Processing, S. (2009) The sequence alignment/map format and SAMtools. Bioinformatics, 25, 2078-2079.

92. Huang, D.W. and Lempicki, R.A. (2009) Systematic and integrative analysis of large gene lists using DAVID bioinformatics resources. Nat. Protoc., 4, 44–57.

93. Love, M.I., Huber, W. and Anders, S. (2014) Moderated estimation of fold change and dispersion for RNA-seq data with DESeq2. Genome Biol., 15, 550.

94. Yin, T., Cook, D. and Lawrence, M. (2012) ggbio: an R package for extending the grammar of graphics for genomic data. Genome Biol., 13, R77.

95. McLean, C.Y., Bristor, D., Hiller, M., Clarke, S.L., Schaar, B.T., Lowe, C.B., Wenger, A.M. and Bejerano, G. (2010) GREAT improves functional interpretation of cis-regulatory regions. Nat. Biotechnol., 28, 495–501.

96. Kent, W.J., Zweig, A.S., Barber, G., Hinrichs, A.S. and Karolchik, D. (2010) BigWig and BigBed: enabling browsing of large distributed datasets. Bioinformatics, 26, 2204–2207.

97. Zhang, Y., Liu, T., Meyer, C.A., Eeckhoute, J., Johnson, D.S., Bernstein, B.E., Nusbaum, C., Myers, R.M., Brown, M., Li, W. et al. (2008) Model-based analysis of ChIP-Seq (MACS). Genome Biol., 9, R137.

98. Liberzon, A., Subramanian, A., Pinchback, R., Thorvaldsdottir, H., Tamayo, P. and Mesirov, J.P. (2011) Molecular signatures database (MSigDB) 3.0. Bioinformatics, 27, 1739–1740.

99. Subramanian, A., Tamayo, P., Mootha, V.K., Mukherjee, S., Ebert, B.L., Gillette, M.A., Paulovich, A., Pomeroy, S.L., Golub, T.R., Lander, E.S. et al. (2005) Gene set enrichment analysis: a knowledge-based approach for interpreting genome-wide expression profiles. Proc. Natl. Acad. Sci. U. S. A., 102, 15545–15550.

100. Shen, L., Shao, N., Liu, X. and Nestler, E. (2014) ngs.plot: Quick mining and visualization of next-generation sequencing data by integrating genomic databases. BMC Genomics, 15, 284.

101. Li, B. and Dewey, C.N. (2011) RSEM: accurate transcript quantification from RNA-Seq data with or without a reference genome. BMC Bioinformatics, 12, 323.

102. Dobin, A., Davis, C.A., Schlesinger, F., Drenkow, J., Zaleski, C., Jha, S., Batut, P., Chaisson, M. and Gingeras, T.R. (2013) STAR: ultrafast universal RNA-seq aligner. Bioinformatics, 29, 15–21.

103. Cong, L., Ran, F.A., Cox, D., Lin, S., Barretto, R., Habib, N., Hsu, P.D., Wu, X., Jiang, W., Marraffini, L.A. et al. (2013) Multiplex genome engineering using CRISPR/Cas systems. Science, 339, 819–823.

104. Ran, F.A., Hsu, P.D., Wright, J., Agarwala, V., Scott, D.A. and Zhang, F. (2013) Genome engineering using the CRISPR-Cas9 system. Nat. Protoc., 8, 2281–2308.

105. Agu, C.A., Soares, F.A., Alderton, A., Patel, M., Ansari, R., Patel, S., Forrest, S., Yang, F., Lineham, J., Vallier, L. et al. (2015) Successful generation of human induced pluripotent stem cell lines from blood samples held at room temperature for up to 48 hr. Stem Cell Rep., 5, 660–671.

106. Kanagaraj, R., Huehn, D., MacKellar, A., Menigatti, M., Zheng, L., Urban, V., Shevelev, I., Greenleaf, A.L. and Janscak, P. (2010) RECQ5 helicase associates with the C-terminal repeat domain of RNA polymerase II during productive elongation phase of transcription. Nucl. Acids Res., 38, 8131–8140.

107. Livak, K.J. and Schmittgen, T.D. (2001) Analysis of relative gene expression data using real-time quantitative PCR and the 2(-Delta Delta C(T)) method. Methods, 25, 402–408.

