## Supplemental Methods for "Integrated Genome and Transcriptome Analyses Reveal the Mechanism of Genome Instability in Ataxia with Oculomotor Apraxia 2"

### Supplementary Methods

#### Quantification and statistical analysis

Statistical details of experiments, including statistical tests, number of events quantified, standard deviation, standard error of the mean, and statistical significance, are reported in the figures and figure legends. GraphPad Prism7 or 8 software (GraphPad) was used for the statistical analyses of cell biology experiments.

#### Analysis of array comparative genomic hybridization data

Data extraction, analysis and visualization were performed using Agilent Cytogenomics 2.7.11.0 software for human samples or the Agilent Feature Extraction 12.0.0.7 and Agilent Genomic Workbench 7.0.4.0 software for mouse samples (Agilent Technologies). Analysis settings were as follows: genome, hg19 (human) or mm9 (mouse); aberration algorithm, ADM-2; threshold: 6.0; window size, 2 kb; aberration filter, ≥ 3 probes and log2ratio ≥ 0.25. The following log2ratio were used to score the aberrations: Amplification, log2ratio ≥ 2; Gain, log2ratio between 0.25 to 2; Loss, log2ratio between -0.25 to -1; Deletion, log2ratio ≤ 1.

#### GREAT analysis

Genomic Regions Enrichment of Annotations Tool (GREAT) was performed as described (McLean et al. 2010).

#### Analysis of gene expression data

Affymetrix gene expression data were analysed using Bioconductor 2.5 (https://www.bioconductor.org) running on R version 2.10.0 (Team 2008). Probeset expression measures were calculated using the Robust Multichip Average (RMA) method with the oligo package (Gautier et al. 2004; Carvalho and Irizarry 2010). Differential gene expression was assessed between AOA2 and control sample groups using an empirical Bayes t-test (limma package) (Smyth 2005). p-values were adjusted for multiple testing correction using the Benjamini-Hochberg method

(Benjamini and Hochberg 1995). Probesets that exhibited an adjusted p-value of 0.05 or less were called differentially expressed. Differentially expressed probes were used to determine pathway and biological process enrichment using the Clarivate Analytics Metacore pathway analysis tool. Pathways or processes that showed FDR < 0.05 were called as enriched.

**Circos plots**

The Bioconductor package ggbio was used to construct the circos plots (Yin et al. 2012). The genomic locations of Aphidicolin Sensitive breakome Regions (ASR), Common Fragile Sites (CFS), Common and Rare Fragile Sites (CRFS), Early Replicating Fragile Sites (ERFS) and Neocarzinostatin Sensitive breakome Regions (NSR) were described previously (Helmrich et al. 2006; Bignell et al., 2010; Fungtammasan et al. 2012; Barlow et al. 2013; Crosetto et al. 2013). The RDC genes were described previously (Wei et al. 2016; Wei et al. 2018).

**Alignment and quantification of mRNA-seq data**

Reads were aligned either against hg19 (human) or mm9 (mouse) and their respective Refseq annotations using STAR v2.5.1b (Dobin et al. 2013) via the transcript quantification software RSEM v1.2.31 (Li and Dewey 2011). The resulting genome alignment BAM files were sorted and indexed using SAMtools 1.3.1 (Li et al. 2009). Duplicate reads were marked using Picard 2.1.1 (http://broadinstitute.github.io/picard). The resulting gene-level estimated read counts were rounded to integers and further analyzed for differential expression using the Bioconductor package DESeq2 1.12.3 (Love et al. 2014).

**Differential expression of mRNA-seq data**

Genes changing between AOA2 and control lymphoblastoid cell lines were tested in paired fashion. Significant genes were thresholded based on a Benjamini-Hochberg FDR ≤ 0.05, absolute fold-change ≥ 1.5 and a minimum normalized read count > 10 in either the AOA2 or control samples. Data were rlog transformed for heatmap visualization, providing variance shrinkage and normalizing with respect to library size.

The heatmap showing 4 AOA2 LCLs was additionally scaled by taking gene-wise z-scores.

Genes changing between human HAP1 WT and Δ*SETX* samples were assessed using an absolute fold change > 1.5, together with a normalized read count > 10 in either the Δ*SETX* or WT sample. The top 10 largest changing genes in each direction were used for heatmap visualization.

Genes changing between mouse *Setx*$^{+/+}$ and *Setx*$^{-/-}$ MEFs were assessed using an absolute fold change > 2, together with a normalized read count > 30. The top 10 largest changing genes in each direction were used for heatmap visualization.

**Gene Set Enrichment Analysis**

Pre-ranked Gene Set Enrichment Analysis (GSEA) analysis was conducted using Bioconductor's "fgsea" package against the Gene Ontology: Biological Process gene collection defined in the Bioconductor package org.Hs.eg.db. Entrez Gene IDs were used for gene-term mappings. Gene sets containing <100 or >500 genes were discarded prior to testing. Genes not associated with an Entrez gene ID were discarded prior to testing. Genes were ranked on the Wald test statistic from the differential expression analysis. Results were thresholded using a Benjamini-Hochberg adjusted pvalue<0.05.

**Overlap test**

Permutation-based overlap tests were performed to assess whether human and mouse CNCs, gains and losses, were enriched for various fragile sites and/or genic regions. Gains and losses were considered independently and the analysis was restricted to the autosomes. Each gains/loss list was permuted 1000 times by assigning random genomic windows in either the hg19 (human) or mm9 (mouse) genome, with sizes equal to the original regions. These permuted windows were not permitted to overlap gaps in the reference genome, overlap each other, or overlap regions in the original gains/losses list. The permutations were then compared to each fragile site list or gene list, and the total number of overlapping base pairs determined. The number of overlapping base pairs between the true regions and the list of interest was compared to the permutation distribution by calculating a z-score and an associated two-tailed p-value.

Overlap tests were also carried out to determine if gains and losses were associated with upregulated and downregulated genes. The locations of differentially regulated genes for both human and mouse were determined by comparison with the hg19 and mm9 NCBI RefSeq databases (Pruitt et al. 2014). For humans, 1250/1310 upregulated genes and 1046/1089 downregulated genes were identified, and for mouse, 666/684 upregulated genes and 458/470 downregulated genes. The locations were then used as the basis for overlap testing versus the gain/loss regions.

**Alignment of ChIP-seq data**

Single-end reads (75 bp) were aligned to the hg19 genome assembly using BWA-MEM 0.7.15 (Li et al. 2009) with default settings. BAM files were sorted and indexed using SAMtools 1.3.1 (Li et al. 2009). Duplicate reads were marked using Picard 2.1.1 (http://broadinstitute.github.io/picard).

**Peak calling**

RNAPII peaks from individual HAP1 WT and Δ*SETX* replicate samples were called against their respective input controls using MACS2 2.1.1 software (Zhang et al. 2008) with the following command line options: '-g hs –q 0.05 –m 5,50'. Peak sets were thresholded for significance based on a q-value ≤ 0.01 and a fold enrichment ≥ 5, before taking the intersect of regions common to both WT or Δ*SETX* biological replicates. Peaks were further restricted to a set within +/- 500 bp of a Refseq gene's TSS.

**Meta-gene profiles**

Meta-gene profiles of RNAPII coverage were created using ngs.plot software (Shen et al. 2014) using standard Ensembl protein-coding gene definitions, n= 20,242. Coverage is represented as Read count Per Million mapped reads (RPM).

**Traveling ratios**

Travelling ratios were calculated as described (Rahl et al. 2010). Briefly, each transcript was divided into: i) a promoter-proximal bin −30 bp to +300 bp around its TSS and ii) a gene body bin to the TTS. The traveling ratio is the ratio of RNAPII density in the promoter-proximal bin to that in the gene body. The most abundant transcript based on mean promoter RPKM across all samples was taken to be representative of the gene and all other transcripts were discarded. Transcripts for which a travel ratio was calculated to be zero (i.e. no reads in the promoter) or infinite (i.e., no reads in the gene body) in any of the samples were removed from the analysis, n = 20,265 transcripts (genes). An additional pre-filter, limiting the analysis to transcripts with a significant RNAPII peak (see peak calling) over their TSS leaving n = 10,132 transcripts (genes).

**Ratio of traveling ratios**

The ratio of traveling ratios (RoTR) was defined as the traveling ratio over a specific transcript for an RNAPII sample, divided by the traveling ratio of its respective control on the same transcript. Transcripts were limited to genes with i) a WT or Δ*SETX* RNAPII promoter peak, ii) ≥ 2kb distant from a neighboring gene on the same strand, and iii) between 2kb and 300kb in width. The most abundant transcript based on mean promoter RPKM across all samples was taken to be representative of the gene and all other transcripts were discarded. For the purposes of visualization, transcripts for which a travel ratio was calculated to be zero (i.e., no reads in promoter) or infinite (i.e., no reads in gene body) in any of the samples were removed from the analysis, leaving 7,492 transcripts (genes). Genes were stratified into width categories based on the quantiles of the gene width distribution: short: <20%, medium: 40-60% and long: >80%. A Wilcoxon rank sum test was used to assess the significance of differences between RoTR distributions.

**GCskew**

GCskew (strand asymmetry in the distribution of guanines and cytosines) was calculated around the TSS (+/- 400 bp) of the 7492 genes identified in the RoTR analysis as follows: GC skew = (G − C)/(G + C)

#### Base composition

The proportion of each base was calculated around the TSS (+/- 400bp or -800 to +400bp) of the 7492 genes identified in the RoTR analysis at single bp resolution. GC-content was calculated as the percentage of the nucleotides that possess either "G" or "C" bases. GCskew was calculated as described. Where appropriate a loess curve was fitted to the data.

#### BigWig files

BigWig files representing genome-wide read depth coverage were generated from BAM alignment files using BEDtools' genomeCoverageBed function (Quinlan and Hall 2010). BedGraph files were in turn converted to bigWig format using the bedGraphToBigWig function from the KentTools package (Kent et al. 2010). Genome browser profiles were generated using the UCSC browser.

#### References

Benjamini Y, Hochberg Y. 1995. Controlling the false discovery rate: A practical and powerful approach to multiple testing. *J. Royal. Stat. Soc. Series B* **57**: 289-300.

Carvalho BS, Irizarry RA. 2010. A framework for oligonucleotide microarray preprocessing. *Bioinformatics* **26**: 2363-2367.

Dobin A, Davis CA, Schlesinger F, Drenkow J, Zaleski C, Jha S, Batut P, Chaisson M, Gingeras TR. 2013. STAR: ultrafast universal RNA-seq aligner. *Bioinformatics* **29**: 15-21.

Gautier L, Cope L, Bolstad BM, Irizarry RA. 2004. Affy-analysis of Affymetrix GeneChip data at the probe level. *Bioinformatics* **20**: 307-315.

Kent WJ, Zweig AS, Barber G, Hinrichs AS, Karolchik D. 2010. BigWig and BigBed: enabling browsing of large distributed datasets. *Bioinformatics* **26**: 2204-2207.

Li B, Dewey CN. 2011. RSEM: accurate transcript quantification from RNA-Seq data with or without a reference genome. *BMC Bioinformatics* **12**: 323.

Li H, Handsaker B, Wysoker A, Fennell T, Ruan J, Homer N, Marth G, Abecasis G, Durbin R, Genome Project Data Processing S. 2009. The sequence alignment/map format and SAMtools. *Bioinformatics* **25**: 2078-2079.

Love MI, Huber W, Anders S. 2014. Moderated estimation of fold change and dispersion for RNA-seq data with DESeq2. *Genome Biol.* **15**: 550.

McLean CY, Bristor D, Hiller M, Clarke SL, Schaar BT, Lowe CB, Wenger AM, Bejerano G. 2010. GREAT improves functional interpretation of cis-regulatory regions. *Nat. Biotechnol.* **28**: 495-501.

Pruitt KD, Brown GR, Hiatt SM, Thibaud-Nissen F, Astashyn A, Ermolaeva O, Farrell CM, Hart J, Landrum MJ, McGarvey KM, Murphy MR, O'Leary NA, Pujar S, Rajput B, Rangwala SH, Riddick LD, Shkeda A, Sun H, Tamez P, Tully RE, Wallin C, Webb D, Weber J, Wu W, DiCuccio M, Kitts P, Maglott DR, Murphy TD, Ostell JM. 2014. RefSeq: an update on mammalian reference sequences. *Nucleic Acids Res.* **42**: D756-D763.

Quinlan AR, Hall IM. 2010. BEDTools: a flexible suite of utilities for comparing genomic features. *Bioinformatics* **26**: 841-842.

Rahl PB, Lin CY, Seila AC, Flynn RA, McCuine S, Burge CB, Sharp PA, Young RA. 2010. c-Myc regulates transcriptional pause release. *Cell* **141**: 432-445.

Shen L, Shao N, Liu X, Nestler E. 2014. ngs.plot: Quick mining and visualization of next-generation sequencing data by integrating genomic databases. *BMC Genomics* **15**: 284.

Smyth GK. 2005. limma: Linear models for microarray data. in *Bioinformatics and Computational Biology Soulutions Using R and Bioconductor. Statistics for Biology and Health* (eds. RC Gentleman, VJ Carey, W Huber, RA Irizarry), pp. 397-420. Springer, New York.

Team RC. 2008. R: A language and environment for statistical computing. in *R Foundation for Statistical Computing*.

Wei PC, Chang AN, Kao J, Du Z, Meyers RM, Alt FW, Schwer B. 2016. Long neural genes harbor recurrent DNA break clusters in neural stem/progenitor cells. *Cell* **164**: 644-655.

Wei PC, Lee CS, Du Z, Schwer B, Zhang Y, Kao J, Zurita J, Alt FW. 2018. Three classes of recurrent DNA break clusters in brain progenitors identified by 3D proximity-based break joining assay. *Proc. Natl. Acad. Sci .U. S. A.* **115**: 1919-1924.

Yin T, Cook D, Lawrence M. 2012. ggbio: an R package for extending the grammar of graphics for genomic data. *Genome Biol.* **13**: R77.

Zhang Y, Liu T, Meyer CA, Eeckhoute J, Johnson DS, Bernstein BE, Nusbaum C, Myers RM, Brown M, Li W, Liu XS. 2008. Model-based analysis of ChIP-Seq (MACS). *Genome Biol.* **9**: R137.
